# Aberrant microglial responses shape hypothalamic circuits in anorexia nervosa

**DOI:** 10.64898/2026.09.18.748216

**Authors:** Jingjing Xu, Emmy Erskine, Barbara Eramo, Karin Zimmer, Chiara Camoglio, Mridul Chaudhary, Tianxu Feng, Funda Orhan, Elisabeth Welch, Lars Selander, Catharina Lavebratt, Tomas Hökfelt, Martin Schalling, Samudyata Samudyata, Carl M. Sellgren, Ida AK Nilsson

## Abstract

Multimodal data indicates that microglia contribute to the pathophysiology of anorexia nervosa (AN). Here, we investigated microglial modulation of hypothalamic circuits – key regulators of energy balance – hypothesising an implication in the prolonged starvation and low body weight of AN. First, we generated microglia as well as hypothalamic and cortical neurons from patient-derived induced pluripotent stem cells (iPSCs) to discover upregulation of synapse-related genes in hypothalamic neurons, reduced microglial uptake of hypothalamic synaptic structures, and a microglial unresponsivness to the satiety hormone glucagon-like peptide-1. In an AN animal model (*anx*/*anx* mouse), spatial transcriptomics indicated hypothalamic microglial activation and disrupted microglia-synapse signaling. Despite an increased microglia density in both the arcuate nucleus (ARC) and the dorsomedial hypothalamus region (DMH), microglia displayed a decreased per cell uptake of synaptic material in Arc. Together, these data suggest that microglial responses shape hypothalamic circuits with possible implications for the maintained negative energy balance of AN.

## Introduction

Anorexia nervosa (AN) is a severe psychiatric disorder characterized by extreme food restriction, intense fear of weight gain, and profoundly distorted body image ^1^. Affecting approximately 1% of the population–with a strong female predominance–AN carries the highest mortality rate among psychiatric disorders, driven by suicide and complications due to extreme starvation ^1,2^. The heritability is estimated around 50-60%, and recent genome-wide association studies highlighted psychiatric as well as metabolic and anthropometric traits, genetically correlated with the disorder, suggesting to repharse AN as a metabo-psychiatric disorder ^3,4^. Psychiatric comorbidities are common, including schizophrenia (SCZ) ^5^ and autism spectrum disorder (ASD) ^6^. A key feature of AN is the long term maintained negative energy balance, failing to activate mechanisms to prevent ‘elective starvation’. This could indicate a failure in the systems regulating energy homeostasis, with disrupted communication between the brain and body. AN-associated genetic variants are also enriched in brain areas associated with food intake and energy homeostasis, including specific hypothalamic regions, as well as on cell level in microglia ^7^. The *anx/anx* mouse, which mimics the core features of AN; starvation and emaciation, display enlarged microglia surrounding degenerating hypothalamic neurons ^8^, further supporting a role of microglia in anorectic conditions. Microglia, primarily known for their rapid response to injury, are also involved in multiple neurodevelopmental processes and in maintaining CNS homeostasis ^9,10^. Deviant microglia-mediated synapse elimination has been suggested as a mechanism of action in other psychiatric disorders^11^ including SCZ ^12^ and ASD ^13^, mentioned above. However, the role of microglial-driven synaptic plasticity in AN remains unclear.

In this study, we first employed an induced pluripotent stem cell (iPSC) derived model of microglial phagocytosis of synapses to discover lower uptake of hypothalamic, but not cortical, synaptic structures in the AN models. In control models, stimulation with the satiety hormone glucagon-like peptide-1 (GLP-1) decreased the uptake of synaptic structures while this effect was not seen in AN derived models. Transcriptomic analysis of AN-microglia revealed downregulation of the GLP-1R and subsequent profiling indicated an antigen-presenting and neurodegenerative microglial profile. We then studied *anx*/*anx* mice and observed a decreased uptake of synaptic structures in microglia within the dorsomedial hypothalamus (DMH), per cell by volume, while the overall total uptake of synapses by microglia cells in the hypothalamus was increased. Spatial transcriptomics suggested altered communication between synapses and microglia with upregulation of microglial activity markers, and profiling of hypothalamic microglia mapped cells to antigen-presenting, phagocytosing and surveillance-like profiles.

## Results

### Immunocytochemical characterization of AN patient-derived microglia and neurons

iPSCs were derived from fibroblasts of 5 AN patients and 6 healthy controls (HC) (Figure 1a) (Supplementary figure 1). The demographic and clinical characteristics of the cohort are summarized in Table 1. While age was not significantly different, BMI at sampling and minimum BMI during adulthood were, as expected, significantly lower in AN compared to the HC group (Table 1). No significant differences in cell yield per experiment were observed between AN patient- and healthy donor-derived microglia (Supplementary Figure 2). Most of the cells displayed a ramified morphology after differentiation and expressed the canonical microglia markers such as Ionized calcium binding adaptor molecule 1 (IBA1) and Purinergic Receptor P2Y12 (P2RY12) (Figure 1b).

**Figure 1.**
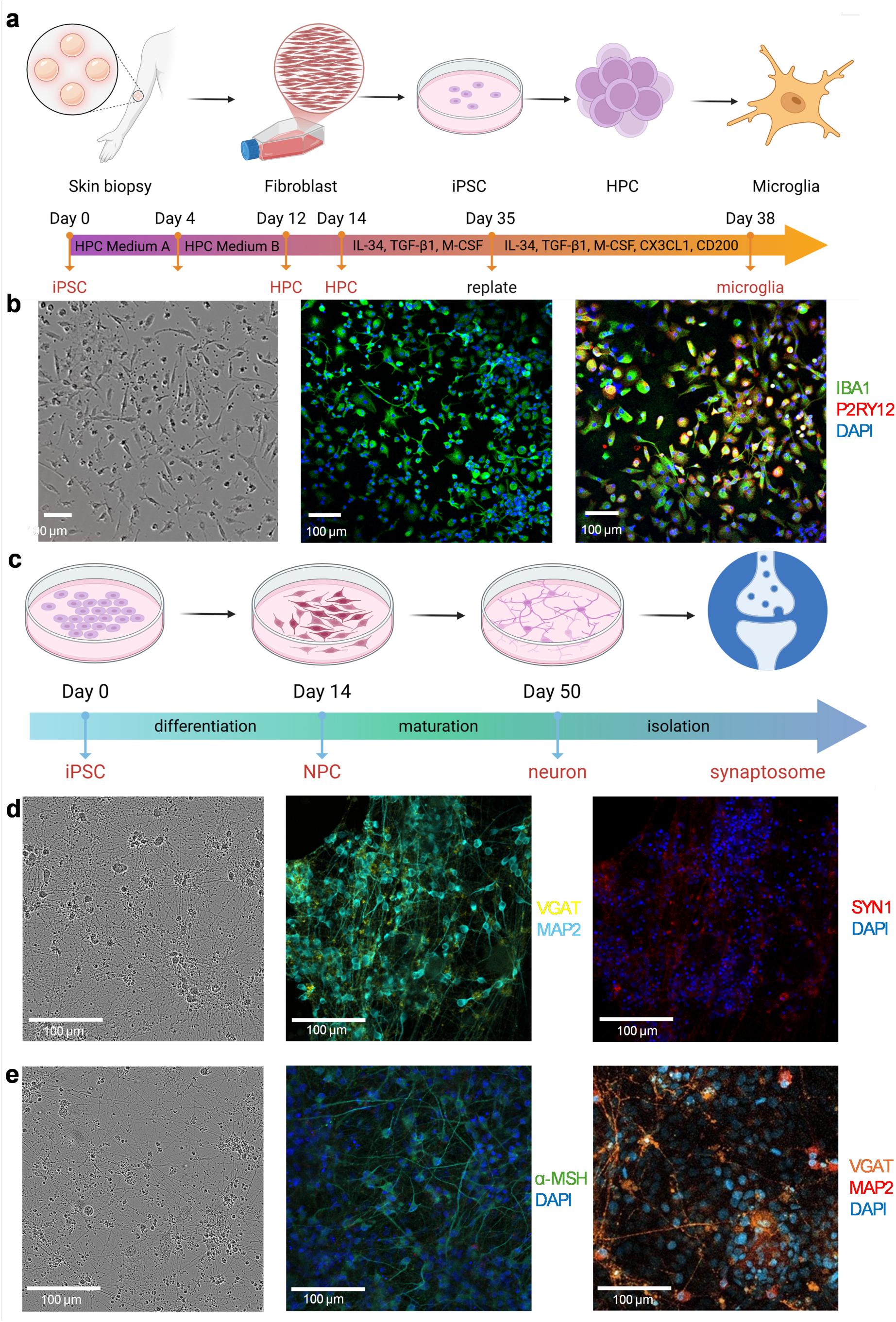
Characterization of AN patient-derived microglia and neurons. **(a)** Outline of the microglia generation protocol, from iPSC induction of skin biopsies to HPCs that differentiate into mature microglia. **(b)** Representative phase-contrast image (Day 38) and immunofluorescence (Day 30 and Day 38) of iPSC-derived microglia with IBA1 (green), P2RY12 (red), and DAPI (blue). **(c)** Outline of the protocol used to generate cortical and hypothalamic neurons, from iPSC induction to NPC that differentiate into mature neurons. **(d)** Representative phase-contrast image and immunofluorescence of iPSC-derived cortical neurons with MAP2 (cyan), SYN1 (red), VGAT (green), and DAPI (blue), **(e)** and of iPSC-derived hypothalamic neurons with α-MSH (green), MAP2 (red), VGAT (orange), and DAPI (blue). HPC, hematopoietic progenitor cell; iPSC, induced pluripotent stem cells; NPC, neural progenitor cells, IBA1, Ionized calcium binding adaptor molecule 1; P2RY12, Purinergic Receptor P2Y12; DAPI, 4’, 6-diamidino-2-phenylindole; MAP2, microtubule-associated protein 2; SYN1, synapsin 1; VGAT, vesicular GABA transporter; α-MSH, alpha melanocyte stimulating hormone. Created in BioRender. Erskine, E. (2026) https://BioRender.com/c93oxb0

**Table 1.** Sex, age, and BMI of the all-female study cohort. Significant differences between AN and HC are marked in bold. AN, anorexia nervosa; BMI, body-mass index; HC, healthy controls; IQR, inter-quartile range.

| Characteristics | AN | HC |
| --- | --- | --- |
| **N** | 5 | 6 |
| **Age at sample (years)** | 34 | 31 |
| **(median [IQR])** | (31-35) | (28-38) |
| **BMI at sample (kg/m<sup>2</sup>)** | **15.0** | **21.2** |
| **(median [IQR])** | **(13.0-15.7)** | **(20.8-22.6)** |
| **Minimum BMI (kg/m<sup>2</sup>)** | **12.4** | **19.0** |
| **(median [IQR])** | **(9.2-12.5)** | **(18.8-20.5)** |

Both cortical and hypothalamic neurons were derived from each iPSC line. By day 50, before extraction of synaptosomes (Figure 1c), both cortical and hypothalamic neurons expressed the pan-neuronal marker (microtubule-associated protein 2) MAP2 and synaptic markers synapsin 1 (SYN1) and vesicular GABA transporter (VGAT) (Figure 1d). Hypothalamic neurons also expressed the anorexigenic neuropeptide α-Melanocyte-Stimulating Hormone (α-MSH) (Figure 1e). No significant differences in yield could be observed between cortical and hypothalamic synaptosomes extracted from AN patients and HC neurons (Supplementary Figure 2).

### Reduced uptake of hypothalamic synaptosomes by AN-derived microglia

Isolated cortical and hypothalamic synaptosomes were labeled with a pH-sensitive dye (pHrodo Red) to indicate localization of synaptic material in the post-phagocytic phagolysosome compartment of microglia. Real-time live imaging was then used to measure the uptake over 18 h (Figure 2a). In AN models, we observed a significantly lower uptake of hypothalamic synaptosomes as compared to HC models (Figure 2b), while no significant differences were observed with regard to uptake of cortical synaptosomes (Figure 2c, Supplementary Table 4).

**Figure 2.**
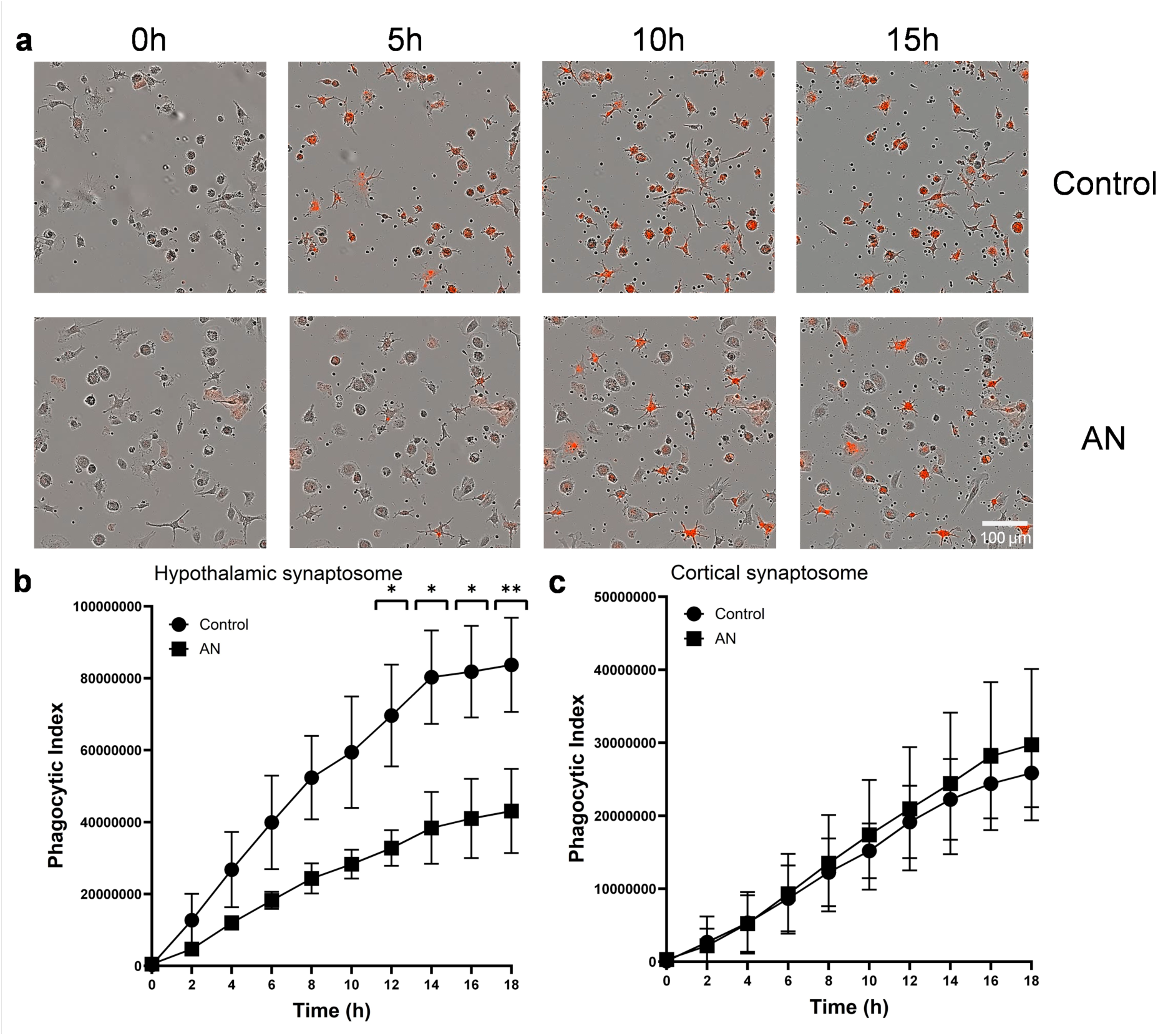
Reduced engulfment of hypothalamic synaptosomes by AN microglia. **(a)** Representative time series images of microglial uptake of pHrodo (red)-labeled hypothalamic synaptosomes during live imaging. **(b, c)** Quantification of the phagocytic index of microglial engulfment of hypothalamic synaptosomes **(b)** and cortical synaptosomes **(c)** during live imaging sessions of 18 h. The phagocytic index is defined as the integrated intensity of pHrodo per well (RCU x µm²/Well) divided by the confluency of the microglia culture (%) and analyzed using two-way repeated ANOVA with Tukey’s multiple comparison tests. Error bars indicate standard error of mean (SEM). AN, anorexia nervosa. * p < 0.05, ** p < 0.01

### Transcriptomic profiling of AN-derived microglia

Bulk RNA-sequencing was performed on AN- and control-derived microglia at baseline and after phagocytosis of cortical and hypothalamic synaptosomes, to further characterize gene expression changes and profile microglia ‘states’, and the transcriptomic profiles were compared. See Supplementary Figure 3 for an overview of differentiations, stimulations and transcriptomic comparisons.

When microglia were at baseline (e.g. not exposed to synaptosomes), we identified 81 differentially expressed genes (DEGs) in the disease model after correcting for multiple testing (Supplementary table 5). This included several upregulated major histocompatibility complex (MHC) class genes such as Human Leukocyte Antigen DQ-alpha 1 and beta 1 in AN (HLA-DQA1 and -B1), typically involved in initiation of the adaptive immune response after processing of debris/pathogens ^14^, thus indicative of an immune activation in AN compared to controls, even in the absence of immune-triggering stimuli. In addition, we observed a downregulation of GLP1R in AN (Figure 3a). Similarly, a non-significant trend towards downregulated GLP1R was also observed in relative expression by qPCR in AN microglia (Supplementary Figure 4).

**Figure 3.**
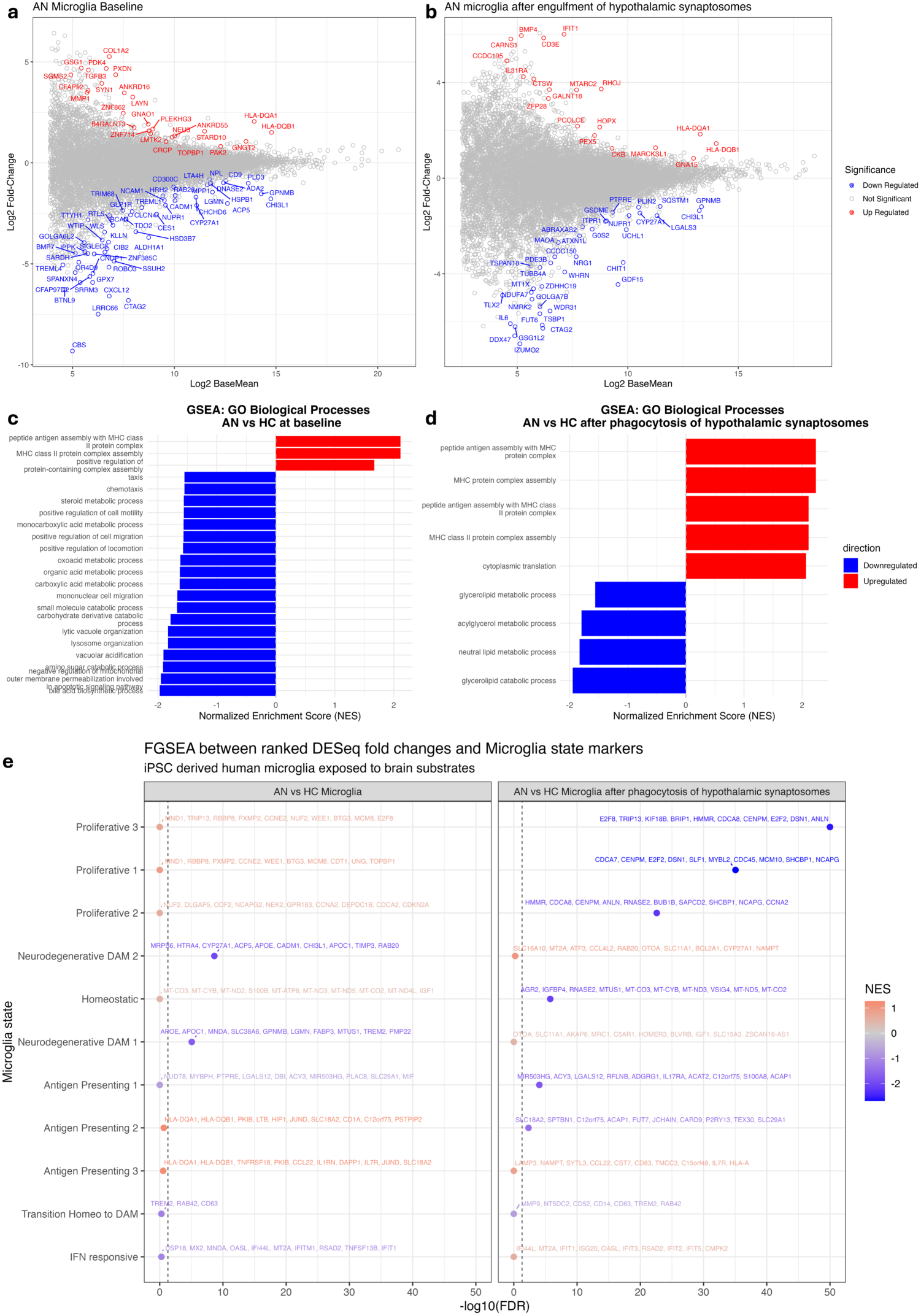
Transcriptomic profile of microglia. **(a)** DEGs of AN patient-derived iPSC-microglia compared to HC at baseline **(b)** DEGs of AN iPSC-microglia after phagocytosis of hypothalamic synaptosomes **(c)** GSEA analysis against GO biological process terms of AN patient-derived iPSC-microglia compared to HC at baseline **(d)** GSEA against GO biological process terms in AN iPSC-microglia after phagocytosis of hypothalamic synaptosomes **(e)** FGSEA between DESeq ranked fold changes against a reference of microglial state markers. DEG; differentially expressed gene; AN, anorexia nervosa; iPSC, induced pluripotent stem cells; HC, healthy control; GSEA, gene set enrichment analysis; GO, gene ontology; FGSEA, fast gene set enrichment analysis.

Gene set enrichment (GSE) analysis of DEGs in AN microglia at baseline was performed, revealing upregulation of antigen presentation and MHC class genes sets, in addition to downregulation of multiple metabolic associated pathways (Figure 3b, d, Supplementary table 6). To gain insight as to the functional state of AN microglia, log fold changes of DEGs were compared against a single-cell iPSC-derived microglial reference datatset exposed to different brain substrates mimicking different disease phenotypes (e.g. amyloid plaques, myelin, synaptosomes) ^15^, using fast gene set enrichment analysis (FGSEA). Downregulated DEGs, in unexposed AN-derived microglia, most closely resembled the neurodegenerative damage associated microglia (DAM) phenotypes, while upregulated DEGs were not significantly associated with any specific phenotype (Supplementary table 7).

In microglia exposed to hypothalamic synaposomes, we identified 57 DEGs, after correcting for multiple testing, in AN compared to control (Figure 3b, Supplementary table 5). Of note, several MHC class genes, including HLA-DQA1 and -B1, were upregulated in both unexposed and phagocytosing AN microglia. Genes downregulated in AN microglia after engulfment of hypothalamic synaptosomes included growth and differentiation factor 15 (GDF15), a stress induced cytokine commonly associated with mitochondrial dysfunction when upregulated ^16,17^ and with a proposed role in energy homeostatic regulation ^18^, but also in microglia engulfment of synapses ^19^. Another downregulated gene of interest in our AN microglia is LGALS3, typically upregulated in neurodegenerative and damage associated microglia (DAM), and involved in initiation of phagocytosis ^20^.

GSEA analysis of DEGs in AN microglia after phagocytosis of hypothalamic synaptosomes indicated downregulation of gene sets associated with lipid metabolic processes (Figure 3d, Supplementary table 6). FGSEA comparing log fold changes of DEGs to the reference datsaset of iPSC derived microglia after exposure to different brain substrates revealed that AN-derived microglia exposed to hypothalamic synaptosomes most closely resembled a profilerative state alongside homeostatic and antigen presenting profiles (Figure 3e, Supplementary table 7).

As previously described, AN microglia displayed lower engulfment of hypothalamic synaptosomes compared to controls, but did not differentially engulf cortical synaptosomes. Thus, AN microglia exposed to hypothalamic synaptosomes were compared to AN microglia exposed to cortical synaptosomes, to characterize the DEGs indicative of the differential microglial engulfment pattern based on origin of synatosomes. We identified 2443 DEGs after multiple corrections in hypothalamic-exposed AN microglia compared to cortical-exposed microglia. GSEA analysis revealed upregulation of mitochondrial associated pathways, and downregulation of multiple pathways associated with the synapse, pre-and post-synaptic signaling and synaptic plasticity (Supplementary figure 5).

As AN-derived microglia had a reduced expression of the GLP1R at baseline, we also assessed DEGs in microglia stimulated with GLP-1. In total, we observed 121 DEGs (Supplementary figure 5) in GLP-1 stimulated AN microglia including maintained upregulation of HLAs, while other immune associated genes were downregulated, such as C-C chemokine receptor 2 (CCR2), a chemokine involved in immune recruitment in several injury and disease states ^21,22^ and long pentraxin 3 (PTX3), an acute inflammatory protein of the pentaxtrin family shared by C-reactive protein (CRP), mediating cross-talk between immune and complement systems^23^.

### AN microglia phagocytosis of hypothalamic synaptosome is unresponsive to GLP-1

To assess the effect of the satiety hormone GLP-1, also having established microglia-modulatory effects^24,25^, on the phagocytic capacity of AN-derived microglia, we pretreated microglia with GLP-1 for 24 h (Figure 4a) prior to adding synaptosomes. In HC derived cells, GLP-1 stimulation significantly reduced the uptake of hypothalamic synaptic structures in microglia, conversely, this difference was not observed in AN derived cells (Figure 4b). On the contrary, when cortical synaptosomes were used, GLP-1 treatment did not influence phagocytic capacity in either HC- or AN-derived microglia (Figure 4c).

**Figure 4.**
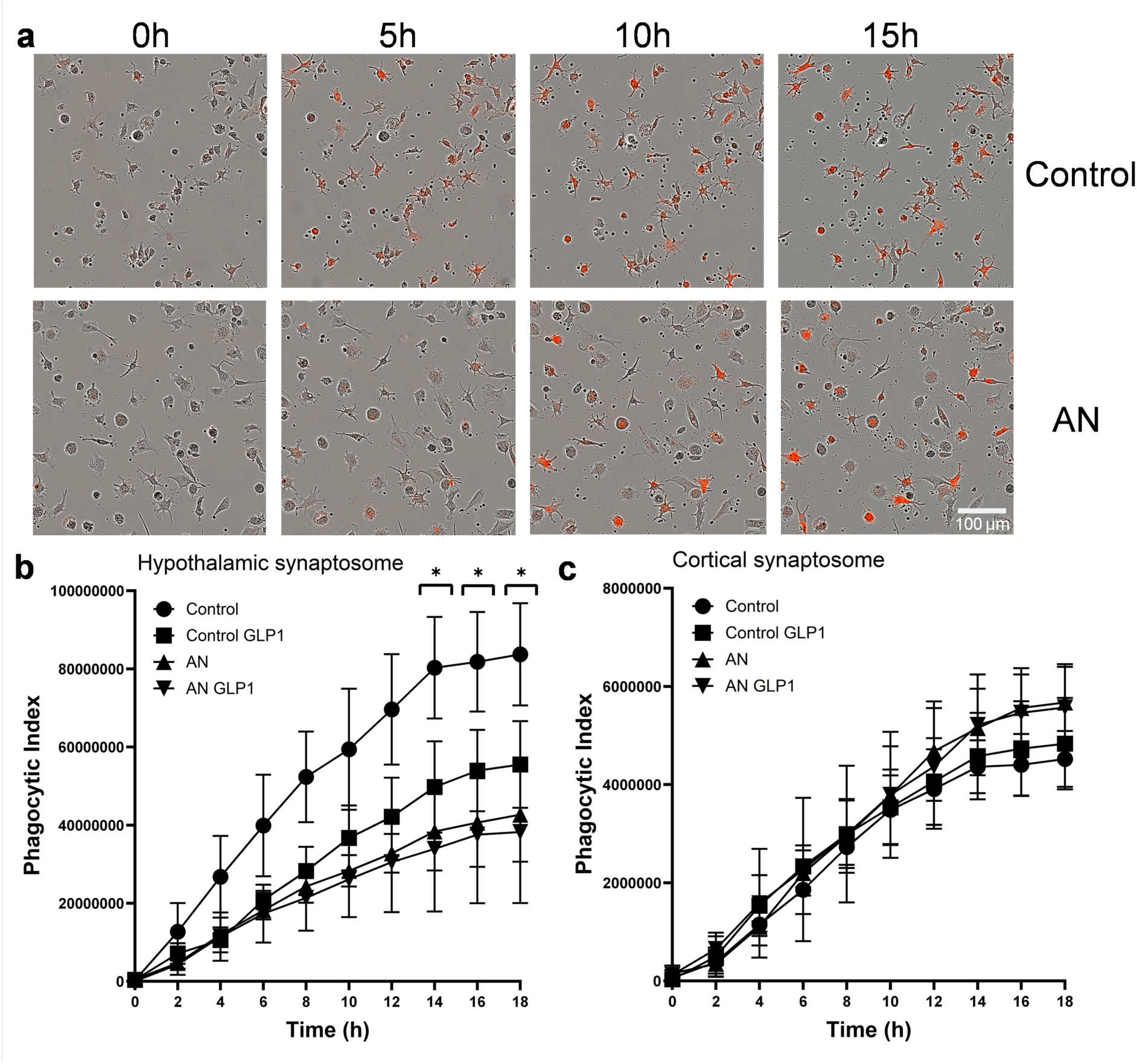
AN microglia phagocytosis of hypothalamic synaptosome is non-responsive to GLP-1 treatment. **(a)** Representative time series images of microglial uptake of pHrodo (red)-labeled hypothalamic synaptosomes upon GLP-1 treatment during live imaging. **(b, c)** Quantification of the phagocytic index of microglial engulfment of **(b)** hypothalamic synaptosomes and **(c)** cortical synaptosomes upon GLP-1 treatment during live imaging sessions of 18 h. The phagocytic index is defined as the integrated intensity of pHrodo per well (RCU x µm²/Well) divided by the confluency of the microglia culture (%) and analyzed using two-way repeated ANOVA with Tukey’s multiple comparison tests. Error bars indicate standard error of mean (SEM). AN, anorexia nervosa; GLP-1, Glucagon-like peptide-1. * p < 0.05, refering to controls with vs without GLP-1 stimulation.

### Transcriptomic profiling of cortical and hypothalamic neurons in AN

Based on the diffrences in microglial engulfment of patient-derived hypothalamic- vs cortical neurons, we performed RNA sequencing of these two neuronal populations. This revealed that the derived hypothalamic neurons exhibit gene expression indicative of a broad hypothalamic profile, expressing e.g., pro-opiomelanocortin (*POMC*), Islet 1 (*ISL1*), nuclear receptor subfamily 5 group A member 2 (*NR5A2*), PR-domain containing member 12 (*PRDM12*), orthopedia homeobox protein (*OTP*), vesicular glutamate transporter 2 (*SLC17A6*/*VGLUT2*), NK2 homeobox 1 (*NKX2-1*) (Supplementary Figure 6), as well as *GLP1R*, leptin receptor (*LEPR*) and insulin receptor (*INSR*). The cortical neurons exhibited gene expression of both glutamatergic and GABAergic neurons with different layer and areal specification, including *SLC17A6*/*VGLUT2*, T-Box Brain Transcription Factor 1 (*TBR1*), SATB Homeobox 2 (*SATB2*), and Reelin (*RELN*) (Supplementary Table 5).

Comparing the transcriptomic profiles of hypothalamic neurons from AN patients versus HCs revealed no significant DEGs after correcting for multiple testing. To nonetheless investigate if non significant genes with large log fold changes impact gene sets, GSE analysis was performed revealing significantly upregulated synapse-related gene sets in AN (Supplementary figure 6). This was particularly interesting in light of the reduced engulfment of hypothalamic synapses seen. In fact, the pathways that ranked the highest enrichment scores were almost exclusively related to the synapse (Supplementary Table 6).

### Spatial transcriptomics of hypothalamic microglia of the anorectic anx/anx mouse

To further evaluate the molecular signature of hypothalamic microglia in an anorectic setting, we performed spatial transcriptomic analysis of 19-21 days old *anx/anx* mice. We selectively profiled IBA1-positive microglia across hypothalamic regions, i.e., the ARC and the DMH (Figure 5a). The DEG profile of the *anx/anx* vs WT microglia included several consistently upregulated genes in ARC and DMH, including beta-2-microglobulin (*B2m)*, complement genes such as *C1qa* and *C1qc,* alongside the inflammatory/immune activation transcripts C-X-C motif chemokine ligand 10 (*Cxcl10)* and Chemokine (C-C motif) ligand 5 (*Ccl5),* and the apolipoprotein *Apoe* (Figure 5b, 5c, Supplementary Figure 9).

**Figure 5:**
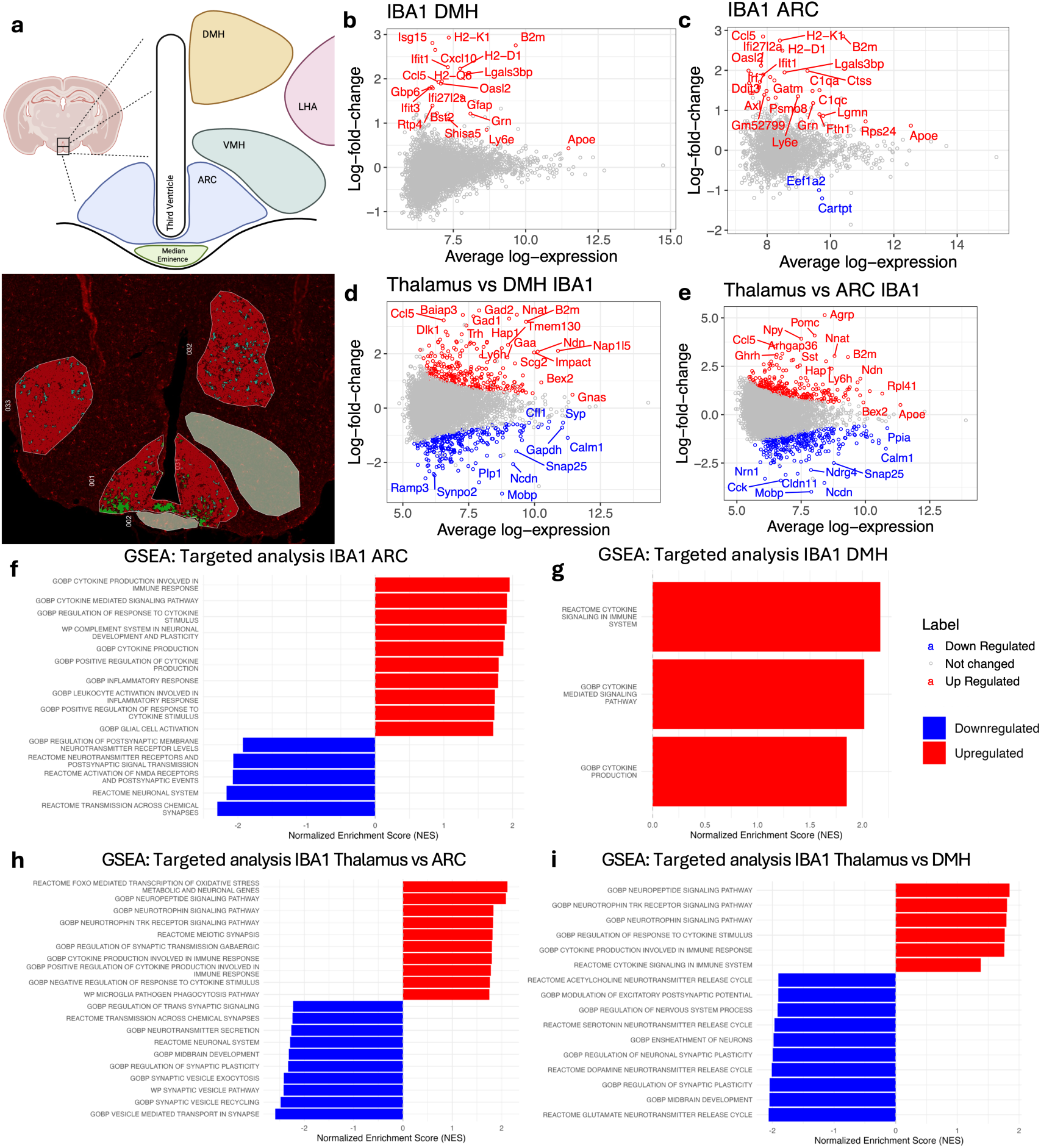
Spatial Transcriptomic analysis of IBA1-positive cells in hypothalamic regions. **(a)** Schematic drawning of hypothalamic regions and GeoMX spatial transcriptomic selection using IBA1-positive signal **(b,c)** DEG analysis of IBA1-positive cells in DMH and ARC, **(d,e)** and thalamus compared to DMH and ARC. Red/blue correspond to upregulated/downregulated genes (p < 0.05) **(f,g)** Targeted GSEA filtered against cell and tissue specific gene set terms in IBA1, ARC, and DMH (**h, i)** Targeted GSEA filtered cell and tissue specific gene set terms of IBA1 in thalamus compared ARC, and thalamus compared to DMH. The colour represents the upregulation (red) and downregulation (blue). IBA1, Ionized calcium-binding adapter molecule 1; DEG; Differentially expressed gene; DMH, dorsomedial hypothalamus; ARC, arcuate nucleus; GSEA, Gene set enrichment analysis; LHA, lateral hypothalamus; VMH, ventromedial hypothalamus. Created in BioRender. Erskine, E. (2026) https://BioRender.com/ia6fr8s

Moreover, GSEA analysis of DEG between anorectic and WT mice was performed. A targeted GSEA approach investigating only gene-sets specific to the microglial cell type and brain tissue highlighted multiple shared upregulated gene sets associated with microglial phagocytosis, the complement system in neuronal development, and inflammatory response. This targeted investigation highlighted a downregulation of gene sets associated with synaptic signal transmission and neuronal systems in the ARC (Figure 5f), not seen in the DMH (Figure 5g), indicative of neuronal ‘contamination’ from synapses/neurons in the close vicinity of the selected microglia being included when segmenting out the microglia population. A broader, untargeted GSEA approach using all GOBP gene-sets indicated differential expression of multiple antigen processing and immune response-associated gene sets. In addition, gene-sets associated with ATP and ribonucleoside triphosphate metabolic effects and the generation of precursor metabolites were affected (Figure 6a, 6b). Network analysis of the significant GOBP gene-sets suggested the involvement of *B2m* as a key component of multiple immune-related pathways in both ARC and DMH, and the interaction of complement components such as *C4b* and *C1q(a-c)* (Supplementary Figure 9).

**Figure 6:**
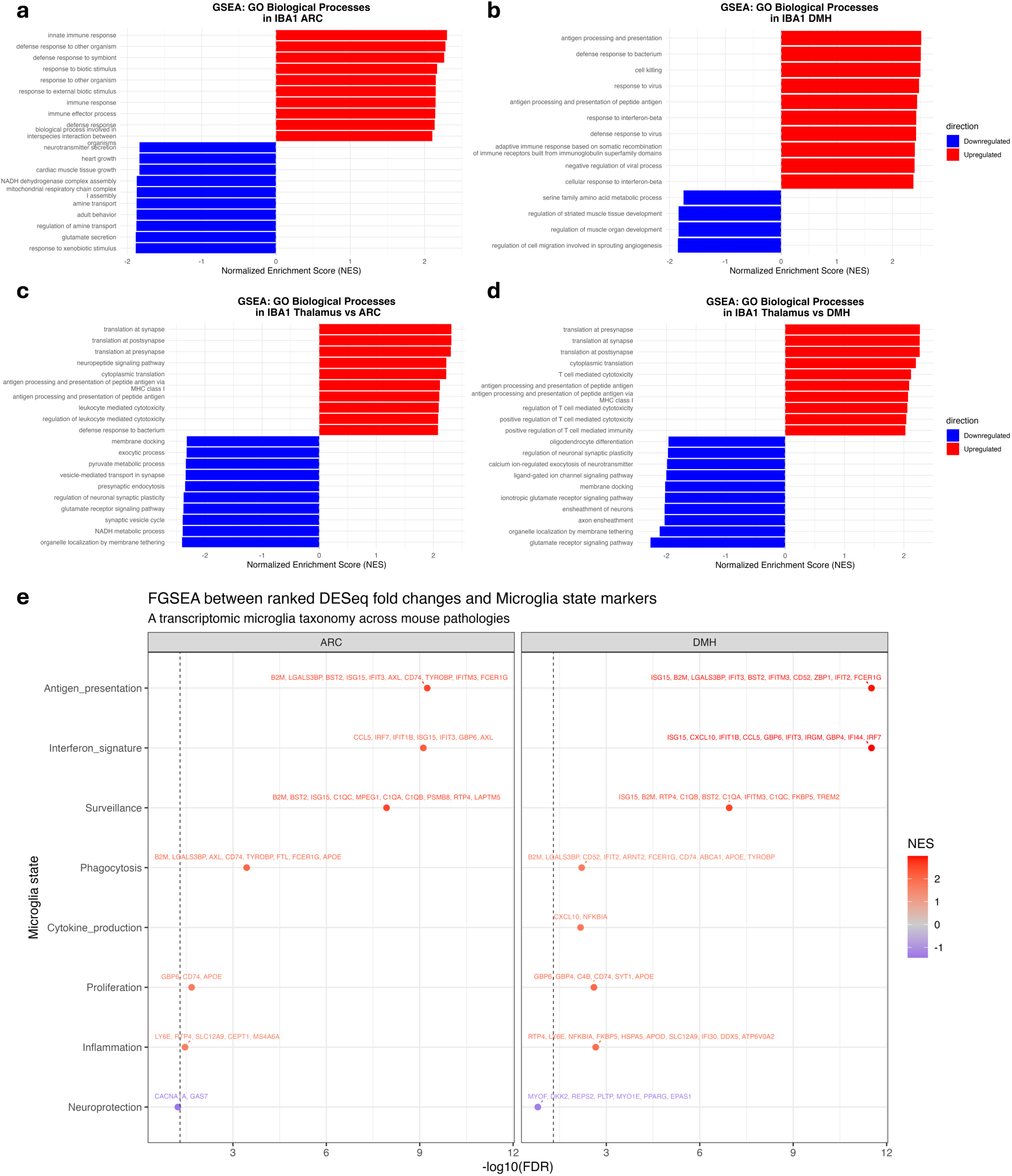
Gene Set Enrichment Analysis GO Biological Processes and Network Analysis of *anx/anx* Spatial Transcriptomics. **(a)** GSEA GOBP of IBA1 in ARC in *anx/anx* compared to WT mice. **(b)** GSEA GOBP of IBA1 in DMH in *anx/anx* compared to WT mice **(c)** GSEA GOBP of IBA1 in thalamus vs ARC in *anx/anx* mice **(d)** GSEA GOBP of IBA1 in thalamus vs DMH in *anx/anx* mice **(e)** FGSEA of DEGs in IBA1 in ARC and DMH in *anx/anx* compared to WT mice against reference atlas of microglia across several neurodegenerative and inflammatory states. GSEA, gene set enrichment analysis; GOBP, gene ontology biological pathway; ARC, arcuate hypothalamus; IBA1, Ionized calcium binding adaptor molecule 1; DMH, dorsomedial hypothalamus; *anx/anx,* anorectic murine model; WT, wildtype; FGSEA, fast gene set enrichment analysis.

To investigate differential molecular signatures between regions with or without enlarged/’activated’ microglia, in *anx/anx* mice, DEG analysis was performed on selected mediobasal hypothalamic regions and the thalamus, respectively, revealing further upregulated immune activation-associated genes in ARC and DMH regions. Some additional indication for neuronal ‘contamination’ as discussed above was observed in the IBA1-positive spatial transcriptomic data, including *Agrp* and *Npy* neuropeptide expression. In both ARC and DMH, synaptosomal-associated protein (*Snap25*) and neurochondrin (*Ncdn*) were downregulated compared to the thalamus, while necdin (*Ndn*) was upregulated in these regions. *B2m* consistently remained upregulated in these regions (Figure 5d, 5e).

Enrichment anaylsis of the enlarged microglia in ARC and DMH compared to microglia in thalamus, of *anx/anx* mice, revealed multiple suppressed synaptic signal transduction and synaptic plasticity-associated gene sets, in line with the DEG findings, as well as upregulated cytokine response and inflammatory pathways when investigating both targeted terms associated to cell type and brain tissue as previously described (Figure 5h, 5i) and untargeted analysis including all GOBP terms (Figure 6c, 6d). Network analysis of the top GSEA terms illustrated suppression of neuronal systems and synaptic plasticity in ARC compared to the thalamus of *anx/anx* mice. Upregulated genes were associated with translation at pre- and post-synapses in the ARC and DMH (Supplementary figure 9).

FGSE analysis was performed to investigate the functional profile of enlarged microglia in the ARC and DMH of the *anx/anx* mouse. Log fold changes of DEGs were compared against a single-cell murine reference atlas comprised of microglia derived from mouse models with a wide range of neurodegenerative and inflammatory disorders ^26^. This analysis revealed that *anx/anx* ARC microglia most closely matched to antigen presentation, interferon signature and surveillance states alongside phagocytic, inflammatory and proliferative profiles. Interestingly, microglia in the DMH matched all these profiles in addition to cytokine production, which could not be assessed in the ARC due to too few gene matches. In fact, the only profile both the ARC and DMH microglia did not significantly resemble was the neuroprotective state (Figure 6e).

### Phagocytosis of hypothalamic synapses by anx/anx microglia

In both the ARC and DMH region of the *anx*/*anx* mouse, we observed an increased microglia density (Figure 7c-d). Performing microglial 3D-reconstructions based on IBA1 staining, we observed internalization of synaptic material as indicated by Syn1/2 expression (Figure 7a-b). Similar to the observations in our patient-derived AN model, microglia in the DMH displayed a significant decrease in synapse uptake per cell in in *anx/anx* mice compared to WTs by volume (Figure 7e), while no significant changes were observed in the ARC (Figure 7f). We observed an overall increased internalization of synaptic puncta by the total pool of IBA-positive microglia in the ARC (Figure 7h) but not DMH (Figure 7g) compared to the total number of synaptic puncta in the region (Supplmentary table 8).

**Figure 7:**
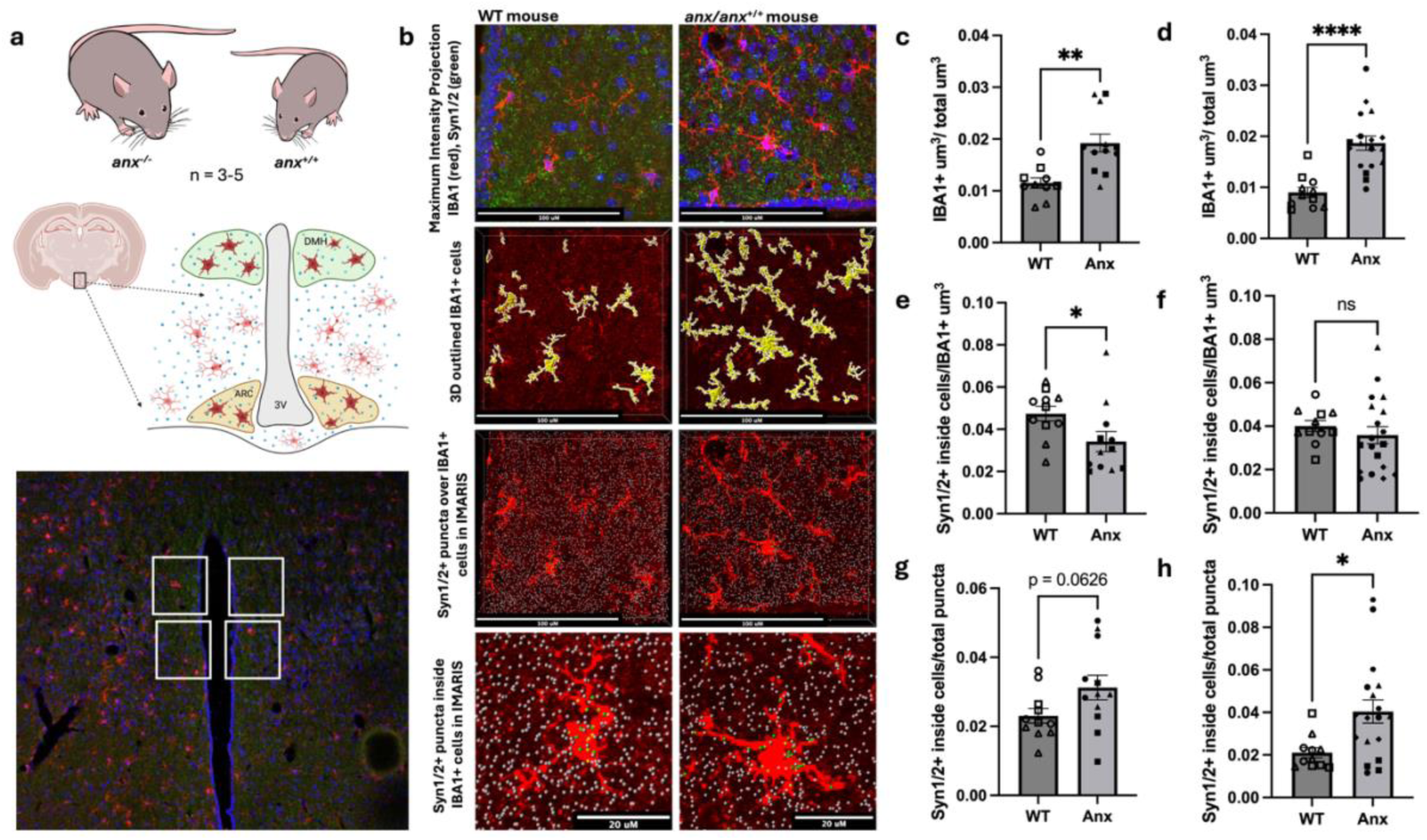
Phagocytosis of ARC and DMH synapses by *anx/anx* microglia. (a) Schematic drawing of *anx/anx* and WT mice and selected ARC and DMH hypothalamic regions. Representative image of DMH with highlighted selected regions, surrounding the third ventricle, imaged at 63X. (b) Figures depict representative Maximum Intensity Projection and IMARIS analysis images of immunofluorescent labelling in the DMH with DAPI (blue), IBA1 (red/yellow), and Syn1/2 (green) in WT and *anx/anx* mice. Quantification of immunofluorescence in DMH (c, e, g) and ARC (d, f, h). IBA1 positive volume (um^3^) across total volume (um^3^) in DMH (c) and ARC (d), internalized Syn1/2 puncta over volume of IBA1 cells (um^3^) in DMH (e) and ARC (f), and number of internalized Syn1/2 spots within the total pool of IBA1 positive microglia over total puncta in DMH (g) and ARC (h). Individual data points represent each image analyzed, and each shape of data points represents an individual animal (3-4 images per animal). Error bars indicate standard error of mean (SEM). WT, wild-type; DMH, dorsomedial hypothalamus; ARC, Arcuate hypothalamus; IHC, immunohistochemistry; DAPI, 4’, 6-diamidino-2-phenylindole; IBA1, Ionized calcium binding adaptor molecule 1; Syn1/2, synapsin; ns, p > 0.05, * p < 0.05, ** p < 0.01, **** p < 0.0001. Created in BioRender. Erskine, E. (2026) https://BioRender.com/2iwbmvf

## Discussion

We observed a reduced microglial uptake of hypothalamic synaptic structures in human AN models and in the *anx/anx* mouse model of AN. In the human models, uptake of cortical synaptic structures was unchanged, while the satiety hormone GLP-1 only decreased microglial uptake of hypothalamic synaptosomes in the control condition. However, complement opsonization proteins such C1q was upregulated in *anx/anx* mice and the total amount of engulfed synaptic material was also increased due to increased microglial volume in the hypothalamic regions. The hypothalamus, specifically the neuronal circuits originating in the mediobasal ARC, are crucial for energy balance, adjusting food intake and metabolism in accordance with the body’s energy status ^27–30^. Despite the extremely low body weight common in AN, patients sustain starvation, often for several years, as though their brains are insensitive to the cues from the body informing them about depletion of energy stores. Thus, we speculate that this reduced phagocytosis of hypothalamic synapses in AN is part of the explanation to the maintained negative energy balance of the disorder, creating a resistance to external hunger cues.

This study is, to the best of our knowledge, the first to report signs of aberrant microglial synaptic engulfment in AN. Previous reports have, however, indicated a role of the complement system in AN, a system with an established role in pruning/engulfment of synapses ^31^. Studies have reported lower *C3* in the serum of patients with AN ^32^, which did not normalize with weight gain ^33^. Other studies reported increasing serum *C3* with increasing BMI ^34^ and parenteral alimentation of AN ^35^. Further genetic association studies of AN, SCZ, and bipolar disorder mapped genes to shared loci harbouring immune-associated genes, such as complement system components *C4A* and *C4B* ^36^. Here complement components are not among our DEGs in AN-derived microglia.

Reduced gray matter volume has been discussed as a proxy for loss of synapses ^37^. On this matter, the largest study to date, in terms of included participants, investigating structural alterations in hypothalamic subregions in AN, reported a regional volume reduction in the hypothalamic region, including ARC ^38^. Hypothalamic focal atrophy ^39^, as well as reduced volume of the total hypothalamus, have also been reported, with reductions in the posterior and inferior tuberal subregions remaining significant after adjusting for total brain volume ^40^. These joint observations may seem contrary to the reduced synapse engulfment seen here. Nevertheless, we cannot with the present design infer the net effect of synapse engulfment by microglia, and thus synapse density and gray matter, in hypothalamus of patients with AN. On the other hand, another study detected no difference in hypothalamic volume in AN ^41^, while a smaller, more recent study reported higher volumes of the global hypothalamus after normalization by the whole-brain volume in individuals with restrictive AN. The same authors reported that the severity of AN was associated with larger structural alterations of several hypothalamic nuclei, including ARC ^42^. Loss of gray matter can also result from inflammation-associated loss of cells, i.e., neurotoxic effect of interleukin 6, as reported in bipolar disorder ^43^. Thus, further studies are needed to establish potential lasting differences in gray matter volume in AN, and if so the mechanism behind this.

Our findings here, indicate a blunted microglial engulfment specifically of hypothalamic synapses, not evident for cortical synapses. In line with this, aberrant hypothalamic connectivity, including increased connectivity in ARC, and feeding-associated alterations of glutamatergic response have been reported in AN ^44^. To further investigate the mechanisms behind this hypothalamus-specific phenomena in AN, RNA-seq was performed comparing iPSC-derived hypothalamic and cortical neuronal populations. Enrichment analysis between iPSC-induced hypothalamic neurons derived from AN patients and HC indicate upregulation of gene sets associated with synaptic membrane (e.g., ‘presynapse’, ‘synaptic membrane’, ‘postsynaptic membrane’), combined with downregulation of structural and ribosomal gene sets. This may indicate basal differences in synaptic connectivity in the AN-derived hypothalamic neurons, which supports the hypothesis that alterations in synaptic plasticity of the hypothalamus may contribute to the pathophysiology of AN.

AN-microglia exhibited consistenly upregulated HLA genes associated with MHC class II, both when unstimulated and after phagocytosis of synaptosomes. HLA facilitates antigen presentation to T-cells, initiating immune responses ^14^, however is also implicated in neuroinflammation in several neurodegenerative, neurodevelopmental and psychiatric disorders ^14,45^. Thus, to investigate the functional profile of AN-microglia compared to controls, FGSE analysis was performed, indicating that the downregulated genes were significantly correlated with neurodegenerative profiles at baseline, while upregulated genes showed no strong tendency toward any microglial profile in particular. Interestingly, after phagocytosis of hypothalamic synaptosomes, microglia matched more closely to proliferative, homeostatic and antigen presenting profiles, this in spite of reduced phagocytosis compared to HC. Adult microglia in the brain self-renew ^46,47^ and respond to apoptotic signals in their microenvironment, proliferating to maintain homeostatic populations ^47,48^. Additionally, reactive proliferation known as microgliosis is a hallmark of many pathological conitions ^49,50^, by which microglia accumulate in regions of dense apoptotic neurons ^50^. This proliferative microglial profile is upregulated in AN while phagocytosis is downregulated, and thus, this proliferative profile may reflect the AN microglial response to the failure of adequately clearing hypothalamic synaptic material. While this analysis cannot inform on whether control-dervied microglia exhibit a homeostatic profile themselves, this analysis does provide some indication into a potentially neurodegenerative-like profile of AN-microglia at baseline. In future, single cell sequencing with cluster analysis would provide greater insight into cellular profiles.

In addition, we report significantly diminished microglial engulfment of hypothalamic synapses after exposure to the satiety hormone GLP-1 in HC-derived microglia, while interestingly no such effect is seen with the AN-derived microglia. The hypothalamic microglial activation seen in high-fat diet (HFD)-induced obese mice can be mitigated by the GLP1R agonist semaglutide ^24^. In fact, GLP-1R agonists have been reported to have ameliorating effects on neuroinflammation in several disease conditions ^51–53^. Moreover, increase in synaptic puncta within the microglial cytoplasm seen with HFD was alleviated during treatment with the agonist, suggesting an effect on phagocytosis in mice ^24^, and semaglutide treatment in murine models of Alzheimer’s disease (AD) has suggested an anti-inflammatory role for the GLP1R-agonist by modulating microglia state ^53^. Interestingly, though our AN microglia continued to exhibit upregulation of antigen presenting genes after GLP-1 stimulation as they did at baseline and phagocytosis, however we also identified downregulation of several immune-associated genes e.g., CCR2 and PTRX3. We speculate that alterations in satiety signaling during treatment with the weight loss drug occur partly through microglia-mediated modulation of synaptic connectivity and plasticity in the hypothalamus. Non-responsiveness of AN-derived microglia to GLP1 in phagocytosis assays may thus be seen as yet another CNS resistance to peripheral signals in AN. Future studies should explore the effect of GLP1 antagonist on AN and HC microglia-mediated phagocytosis.

Further supporting this hypothesis, sequencing of patient-derived microglia reveals DEGs in microglia of AN patients, including a reduction of *GLP1R.* Similarly, a qPCR of GLP1R in which AN microglia illustrated a trending reduction in the relative expression of the receptor compared to controls. This could explain that we see a reduced engulfmen of synaptosome in HC but not AN derived microglia after GLP1 exposure. Taken together these data illustrate aberrances in GLP1 metabolic signaling in AN patient-derived microglia.

The aberrant role of microglial engulfment of hypothalamic synapses is further supported in the *anx/anx* mouse, a rodent model mimicking key aspects of AN, including starvation and maintained negative energy balance ^54^. Multiple regions of the *anx/anx* hypothalamus displayed microglial branching and enlarged cell bodies, as has been reported previously ^8,55^. Here, we report DEGs in hypothalamic microglia, including upregulated complement system genes, e.g., *C1q(a-c)*. In addition to upregulated MHC class-II observed in the human model, several MHC class I-related molecules, e.g*., B2m, H2-K1, and H2-D1*, were upregulated in *anx/anx* hypothalamic microglia in the ARC and DMH, in line with our previous reports ^55^. As already mentioned, complement components are implicated in synaptic engulfment/pruning during healthy development and also in the pathophysiology of other major psychiatric conditions ^56,57,58^. *C1qb* mediates classical microglial phagocytosis pathways, tagging synapses, initiating downstream targeting of synapses for internalization ^59^. MHC class I molecules have also been associated with synaptic plasticity, among others, by modulating the interactions between microglia and synapses ^60–62^.

Moreover, differences in *anx/anx* microglia between hypothalamic and thalamic regions, i.e., regions with and without enlarged microglia, revealed upregulation of complement system genes, including *C4b*, *C1q(a-c),* and MHC class I-associated genes. Consequently, these results support microglial activity in the mediobasal hypothalamus of the *anx/anx* mouse being related to the complement system and synapse remodeling, and illustrate a specificity of these complement components to regions with enlarged microglia. Additionally, APOE is significantly upregulated in the ARC of the *anx/anx* mouse. Microglial APOE is associated with inflammatory response and metabolic dysfunction in AD pathology and neuroinflammation ^63,64^. The APOE4 isoform expressed in microglia was reported to have increased phagocytic capacity *in vitro* ^64^, and APOE4 induced a pro-inflammatory state in human microglia ^65^. Thus, the aberrances in APOE gene expression in our AN murine model may warrant further investigation.

3D visualization of microglia and synaptic puncta in the hypothalamus of the *anx/anx* mouse reveal deviant internalization of synapses in line with the human *in vitro* data. The significant decrease of synaptic internalization by microglial volume in the DMH of the anorectic mouse is in line with the reduction in hypothalamic-specific phagocytic capacity in AN-derived microglia. Nonetheless, we observe an increase in total synaptic uptake in all microglia in the ARC of *anx/anx* compared to WT mice, with a similar trend in the DMH. Taken together with the human data, these results support aberrant microglial phagocytosis in the hypothalamus with significant decreases in synapse-phagocytosis by individual microglial cells in both human and murine models. However, in spite of this blunted phagocytic capacity, the overall increase in microglial cell number and microglial volume due to cell enlargement may cumulatively compensate for this individually reduced effect, explaining the greater overall internalization. We have however, as already mentioned, not investigated total synapse consumption by microglia in AN-patients, and can only speculate as to the total synapse internalization in the patient hypothalamus. These results are in line with the spatial transcriptomic data, indicative of overall increased phagocytic activity and reduction in synapse-associated pathways. Further quantification of synapse maturity and structure could be performed to continue the characterization of the dynamics of microglial synaptic phagocytosis in AN, alongside repeating experiments with a larger number of animals to increase power.

In further support of this aberrant phagocytic role in AN, enrichment analysis comparing DEGs between *anx/anx* and WT mice against murine microglial profiles in a reference dataset profiled *anx/anx* microglia in the ARC and DMH to interferon responsive, antigen presenting, inflammatory, phagocytic, proliferative and surveillance phenotypes, and additionally, DMH microglia matched cytokine-producing profiles. Strikingly, microglia in neither regions resembled the neuroprotective-like profile. While a neurodegenerative profile was not present in the reference dataset, upregulation of interferon pathways is commonly associated with neurodegeneration ^66,67^, which, together with proliferation and antigen presentation resembles the profiling of patient-derived iPSC-microglia which exhibited neurodegenerative profiles at baseline, alongside proliferation and antigen presentation after phagocytosis of hypothalamic synaptic material. Taken together, these results support the hypothesis of aberrances of microglial phagocytic and inflammatory processes in AN.

Finally, we note that *GDF-15* is downregulated in AN-derived microglia after phagocytosis of hypothalamic synaptosomes compared to HC. Intrerestingly, this stress induced cytokine mediates aversive dietary response and has been implicated in both anorexia/cachexia ^68^ and obesity as a marker of nutritional stress ^18^. Interestingly, in a murine model of sepsis-induced cognitive impairment, GDF15 levels were elevated after lipopolysaccharide (LPS) injections via the NF *κ* B pathway, however inhibition of GDF15 alleviated microglial activation and synapse engulfment and synapse-loss, mitigating cognitive and memory-symptoms ^19^. Additionally, in AN, previous studies in lab have identified subgroups of AN patients with elevated GDF15 levels in plasma ^69^ though its role in AN microglia remains uncertain.

Taken together, our data from patient-derived iPSCs and animal model suggest that a reduced microglia-driven synaptic engulfment of hypothalamic neurons in AN is a potential explanation to the maintained negative energy balance and seemingly resistance to peripheral hunger signals of the disorder.

## Materials and methods

### Study individuals

Individuals with primarily restrictive AN according to DSM-IV or V criteria for AN ^70,71^ were informed about the study by clinicians at a specialized eating disorder clinic in Stockholm, Sweden. Further inclusion criteria were female sex, at least 18 years of age, meeting a sickness duration of at least five years, and a maximum of five episodes of purging behavior during the illness. Healthy controls (HC) were recruited via advertisements on social media and locally at ki.se. These age-matched normal-weight female controls reported no personal or family history (parents or siblings) of disordered eating behaviour or autism spectrum disorder. The study was approved by the Regional Ethics Review Board in Stockholm. All participants provided written informed consent.

### Skin sampling and fibroblast culture

Following subcutaneous exposure to a local anesthetic, a 3.0 mm punch tool was used to obtain a dermal biopsy from the nondominant forearm. Biopsies were collected into DMEM GlutaMAX™ medium (Gibco, Waltham, MA, USA) with 20% fetal bovine serum (FBS) (Gibco) and 1% penicillin-streptomycin (P/S) (Gibco). After washes in phosphate-buffered saline (PBS), biopsies were cut into small pieces, transferred to a Petri dish with the dermal side down, and placed in an incubator at 37 °C for 15 min. DMEM GlutaMAX™ (Gibco) with 20% FBS and 1% P/S was gently added to the pieces of skin and changed twice a week until the fibroblasts covered the entire plate. Fibroblast cultures were passaged twice and tested for mycoplasma using MycoAlert™ Mycoplasma Detection Kit (Lonza, Basel-Stadt, Switzerland) before the cells were collected for freezing and reprogramming.

### Induced Pluripotent Stem Cell reprogramming

Human fibroblasts were reprogrammed, and the resulting iPSC colonies stabilized and expanded under xeno-free conditions by BrainXell (Madison, WI, USA). Each line was plated into six-well plates without feeders at three different plating densities and subjected to messenger RNA reprogramming. Colonies were bulk passaged from the most productive well to establish passage #1 iPSC cultures on iMatrix-511 SILK (Nippi, Tokyo, Japan) in Essential 8™ medium (Gibco) and expanded in the same culture system until at least passage #3 before frozen down for storage.

### Karyotyping

iPSC cultures were tested for mycoplasma as described above, after which the cells were collected for genomic DNA extraction using DNeasy Blood & Tissue Kit (Qiagen, Venlo, Netherlands). Genomic DNA was sent to Life & Brain GmbH (Bonn, Germany) for a low-resolution karyotyping based on genotyping using an Illumina microarray. Karyotyping revealed copy number variations larger than 500,000 in two control iPSC lines (Supplementary Table 1). Nevertheless, both lines exhibited typical pluripotency markers and retained self-renewal and differentiation capacity, and thus were included in the study (Supplementary Figure 1).

### Microglia differentiation

Differentiation of iPSCs to microglia was performed according to a published protocol ^72^ with minor changes (Figure 1a). Small clusters of iPSCs were differentiated into hematopoietic progenitor cells (HPCs) using the STEMDiff hematopoietic kit (STEMCELL Technologies, BC, Canada). HPCs harvested from day 10 to day 14 were replated on 1% Geltrex™ (Gibco)-coated 6-well plates and cultured for another 25 days in a differentiation medium DMEM GlutaMAX™ (Gibco), 2x insulin-transferrin-selenite (Gibco), 1x MEM Nonessential Amino Acids (Gibco), supplemented with 2x B27, 0.5x N2 (Gibco), 400 μM monothioglycerol (Sigma-Aldrich, MO, USA), 5 μg/mL insulin (Sigma-Aldrich), 100 ng/mL interleukin 34 (PeproTech, Rocky Hill, USA), 50 ng/mL transforming growth factor-β-1 (PeproTech), and 25 ng/mL macrophage colony-stimulating factor (PeproTech). The medium, including fresh cytokines, was changed every other day. 100 ng/mL CD200 (PeproTech) and 100 ng/mL CX3CL1 (PeproTech) were added for the last 3 days to enhance microglia maturation.

### Neuronal differentiation

Hypothalamic and cortical neurons were generated according to a published protocol ^73^ with minor changes (Figure 2a). In brief, iPSCs were maintained in StemFlex™ medium (Gibco) for at least two passages on 1% Geltrex-coated 6-well plates before differentiation started. Differentiation medium consisted of a 1:1 mixture of Neurobasal (Gibco) and DMEM GlutaMAX™(Gibco), supplemented with 1x N2 (Gibco), 1x B27 (Gibco), 1% P/S (Gibco), 1% sodium bicarbonate (Gibco), and 1% MEM Nonessential Amino Acids (Gibco). XAV939 (Tocris, Bristol, UK), LDN-193189, and SB431542 (Sigma-Aldrich) were added at decreasing concentrations from day 0 to day 10. For the hypothalamic protocol, 1 μM ventralizing factors, smoothened agonist, and purmorphamine (Sigma-Aldrich) were added from day 2 to day 8, and 5 μM DAPT (Sigma-Aldrich) from day 8 to day 14 to promote neurogenesis. For the cortical protocol, fibroblast growth factor 2 (Sigma-Aldrich) was added from day 12 to day 14. Day 14 neural progenitor cells were replated onto 2% Geltrex-coated 6-well plates and maintained in differentiation medium supplemented with 2 µM PD0332991 (Sigma-Aldrich), 1 µM LM22A4, 300 µM γ-aminobutyric acid (GABA), 2 µM CHIR99021 (Tocris), and 10 ng/ml brain-derived neurotrophic factor (PeproTech) from day 16 to 50. 4 µM DAPT and 1 µM NHK477 (Sigma-Aldrich) were added from day 16 to 21. The medium was changed every other day.

### Immunocytochemistry

Day-30 microglia and neurons were replated on black 96-well optically clear bottom plates and cultured until maturation. Day-38 microglia and day-50 neurons were fixed in 4% paraformaldehyde (PFA) (Thermo Scientific), permeabilized for 5 min in 0.3% Triton™ X-100 (Sigma) in PBS, and blocked for 1 h at room temperature (RT) in 5% donkey serum in PBS. Primary antibodies were diluted in 1% serum in PBS and incubated with cells at 4°C. The following day, the cells were washed and incubated with secondary antibodies (1:500) for 1 h at RT, washed twice with 0.3% Triton X-100 in PBS, before being incubated with 4,6-diamidino-2-phenylindole, dihydrochloride (DAPI) (Invitrogen) (1:500) for 5 min. Microglia were immunolabelled with antisera against IBA1 and P2RY12, hypothalamic neurons with antisera against α-MSH, MAP2, VGAT, and DAPI, and cortical neurons with antisera against SYN1, MAP2, VGAT, and DAPI. A full list of antisera used can be found in Supplementary Table 2.

### Synapse preparations

Day-50 neurons cultured in 6-well plates were collected in Syn-PER™ Synaptic Protein Extraction Reagent (Thermo Scientific, Carlsbad, USA) to extract synaptosomes according to the manufacturer’s instructions. Concentrations of the synaptosomes were determined by Pierce BCA Protein Assay Kit (Thermo Scientific) before being stored in 5% dimethyl sulfoxide (Sigma-Aldrich) at -80°C.

### Phagocytotic assay

Day-30 microglia were replated at 3,000 cells per well, three wells per condition, on black 96-well optically clear bottom plates (Thermo Scientific) and cultured until day 38, with or without stimulation with 100 ng/ml glucagon-like peptide-1 (GLP-1) for 24 h. Thawed synaptosomes were dissolved in 0.1M sodium bicarbonate buffer and labeled with pHrodo™ Red (Invitrogen, Carlsbad, CA, USA) (1 μg per 20 μg synaptosome) at 4°C for 2 h. The homogenized solution was washed by adding sodium bicarbonate buffer and centrifugation at 15,000g for 15 min. Pelleted synaptosomes were resuspended in PBS and sonicated for 30 min. Day-39 microglia were fed with 2 μg synaptosomes per well, and the plate underwent live imaging using Incucyte S3 Live-Cell Analysis System (Sartorius, Göttingen, Germany) for 18 h. Whole well images were taken every hour, and both the phase and red channel were analysed using the Incucyte software (version 2022A). For stimulations with cortical synaptosomes, three and four biological replicates were included in HC- and AN-derived microglia respectively. For stimulations with hypothalamic synaptosomes, three and five biological replicates were included in HC- and AN-derived microglia respectively. For stimulations with GLP-1 in combination with phagocytosis of hypothalamic synaptosomes, three and four biological replicates were included in HC- and AN-derived microglia respectively. For stimulations with GLP-1 in combination with phagocytosis of cortical synaptosomes, four biological replicates were included both groups.

### RNA sequencing

Day-30 microglia and neurons were replated on 96-well plates, with two wells per line and three lines, and cultured until day 38 and day 50, respectively. As replicates were plated and grown in individual wells, and separate libraries were prepared and sequenced per replicate, these were considered biological replicates. Thus, six replicates of AN-derived microglia and six replicates of HC-derived microglia were sequenced. Cell culture medium was aspirated, and the plates were stored at -80°C before library preparation. cDNA libraries were generated using the Smart-seq3 protocol ^74^, and sequenced using the NovaSeq X Sequencing System (Illumina, San Diego, California, USA).

### qPCR

Cells were collected in 400 μL Buffer RLT included in the RNeasy Plus Micro Kit (Qiagen, Venlo, Netherlands), and RNA was extracted according to the manufacturer’s instructions. cDNA was synthesized from RNA using SuperScript III First-Strand Synthesis System (Invitrogen) according to the manufacturer’s instructions. qPCR was performed in three replicates using iTaq™ Universal SYBR Green PCR kit (Bio-Rad, Hercules, USA) on a QuantStudio™ 6 Real-Time PCR Instrument (ThermoFisher, Waltham, USA). The PCR reaction consists of 95°C for 10 min, 95°C for 15 sec, and 60°C for 60 sec for 40 cycles, and 72°C for 5 min. The measured transcript abundance of GLP1R was normalized to beta-actin (ACTB) using the delta-delta cycle threshold method. Primer sequences can be found in Supplementary Table 3.

### Animals

The regional animal ethics committee of north of Stockholm approved all animal experiments. Heterozygous *anx* breeding pairs (B6C3Fe–a/a–anxA/+ a), initially obtained from the Jackson Laboratory (Bar Harbor, ME), were used to set up an intercross. Genotyping was performed using simple sequence length polymorphism markers mapped to the sub-chromosomal region, where the anx mutation is located ^75,76^. The *anx/anx* mice and their wild-type (WT) siblings were housed at RT (22°C) with a 12:12-h light-dark cycle and free access to food and water, i.e., milk from the mothers. Mice were perfused via the ascending aorta with Tyrode’s Ca^2^-free solution at 37°C, followed by fixation with a mixture of 4% PFA in 0.16 M phosphate buffer (pH 6.9 at 37°C) and then with the same ice-cold fixative at postnatal day 19-21. Brains were rapidly dissected out and snap frozen. Coronal brain sections (10 μm), including the hypothalamus, were cut on a cryostat (Cryostar NX70, Thermo Fischer Scientific, Walldorf, Germany) and thaw-mounted on Superfrost Plus Slides (VWR, Leicestershire, UK).

### Spatial transcriptomics

For spatial transcriptomics, four animals per genotype (one to two sections per animal) were included in analysis of ARC and DMH, while three *anx/anx* mice were included in analysis of thalamus, with average weights (±SD) of 4.8±1.4 and 8.4±0.4 g (P<0.01) for *anx/anx* and WT mice, respectively. Tissue slides were prepared in accordance with the Manual Slide Preparation Guide from Nanostring (MAN-10150). In brief, slides were fixed for 4 h in 10% neutral buffered formalin (Sigma-Aldrich), followed by washing in PBS and baking at 60°C for 30 min. Subsequently, the tissue slides were dehydrated by consecutive immersions in 50%, 70%, and 100% ethanol, 5 min each. Antigen retrieval was performed using Tris-EDTA buffer (pH 9) (Invitrogen) at 85°C for 15 min, followed by one wash in PBS, after which the tissue slides were incubated with antisera against IBA1 (1:200) at 4°C. The following day, Alexa 647-conjugated secondary antibody (1:500) was added to slides. To expose RNA targets, tissue slides were subjected to mild digestion using 0.1µg/mL Proteinase K (Invitrogen) at 37°C for 15 min. Hybridization of the RNA probes (Mouse Whole Transcriptome Atlas from NanoString, Seattle, Washington, USA) was conducted at 37°C. Stringent wash performed in a mix of 50% 4X SSC (Sigma-Aldrich) and 50% Formamide (Sigma-Aldrich), twice at 37°C for 25 min, followed by two washes in 2X SSC (Sigma-Aldrich). The tissue slides were incubated with nuclear stain SYTO 83 (1:10 in TBS-T, Nanostring) for 15 min, rinsed in TBS-T, and loaded onto the GeoMx DSP. Microglia cells were identified via the fluorescent signal from the IBA1 antiserum. By selectively illuminating these cells with UV light, the photocleavable DNA oligos identifying each transcript were released, aspirated, and subsequently identified and ‘counted’ using a NovaSeq SP-100 v1.5 flow cell (Illumina).

### Immunohistochemistry

Immunohistochemistry (IHC) was performed on hypothalamic tissue. In the analysis of ARC and DMH regions five and three *anx/anx* mice were included in each comparison respectively, and three WT mice, with average weights of (±SD) 5.4±1.0 and 7.6±1.4 g (p < 0.09) for *anx/anx* and WT mice, respectively. Slide-mounted sections were used, using sequential staining with tyramide signal amplification (TSA) for signal amplification IHC ^77–80^, followed by conventional IHC^8,81^. Sections were incubated with SYN1/2 rabbit antiserum (overnight (O/N), 4°C, 1:3,000) followed by anti-rabbit immunoglobulins conjugated to HRP (Agilent Technologies, 1:200). Thereafter, slides were incubated with tyrosine residues conjugated with FITC (AKOYA Biosciences, 1:200) in Amplification Diluent (1X, AKOYA Biosciences), all in accordance with the manufacturer’s instructions. Thereafter, heat-mediated citric acid antigen retrieval (90°C, 5 min) was performed, and slides were incubated with IBA1 rabbit antiserum (4°C O/N, 1:200), before incubation with Alexa Fluor 555 donkey anti-rabbit (30 min, 1:100) (Thermo Fisher Scientific). Slides were mounted with Glycerol with DAPI and 1,4-diazabicyclo 2.2.2 octane (DABCO, Abcam). Antisera are listed in Supplementary Table 2.

### Confocal Microscopy

Images were acquired on a Zeiss LSM 880 confocal microscope (Carl Zeiss, Jena, Germany) operated with ZEN software (Black edition). Z-stacks were collected at 1μm intervals, using a 63x oil-immersion objective. Images for both brain regions of interest, ARC and DMH, were acquired from the central portion of each structure using the third ventricle as an anatomical landmark.

### Image Analysis

Three-dimensional reconstruction and quantification were performed using IMARIS 10.2.0 software (Bitplane, Zurich, Switzerland). In each group 3-4 images were acquired per animal and per region. TIFF stack images were converted to IMS format using the Imaris file converter, and voxel dimensions (X: 0.095 μm, Y: 0.095 μm, Z: 1 μm) were manually entered based on the confocal acquisition metadata. Analysis was carried out for each channel separately. Syn1/2 immunofluorescence was quantified using spot detection with an estimated XY diameter of 0.095 μm. The IBA1 immunofluorescence signal was quantified using the cell tool to segment cells displaying prominent cell bodies. The segment tool gave an estimate of the total volume of fluorescence in each image, used to normalize volume-based reads against the total 3D volume of each section. Finally, the cell module was used to overlay spot and cell signal, enabling the quantification of synapses within and outside microglia.

### Statistical analysis and transcriptomic analyses

Statistical analysis of phagocytosis assay, qPCR, and IHC data, including co-stained IBA1 and Syn1/2, analyzed using IMARIS, was performed in GraphPad Prism 10. For the phagocytosis assay, the phagocytic indexes from all timepoints were automatically calculated by the Incucyte software, defined as the integrated intensity of pHrodo per well (RCU x µm²/Well) divided by the confluency of the microglia culture (%). The data were then analyzed using a two-way repeated ANOVA with Tukey’s multiple comparison tests. For the IMARIS data, normality of distribution was tested using Shapiro-Wilk test alongside Grubbs outlier test (alpha = 0.05). Samples following normality were tested using an unpaired two-tailed t-test, and non-normal samples were tested with the Mann-Whitney two-tailed test. Significance was determined as a p-value of <0.05.

For RNA sequencing data, raw demultiplexed FASTQ files were merged and processed with the nf-core/rnaseq pipeline ^82^, and reads were mapped against GRCh38 and quantified with gene annotations from GRCh38.113. STAR (2.7.11b) ^83^ for alignment. Principal component analysis (PCA) was carried out based on log2-normalized variance-stabilized transformed counts to identify and remove sample outliers. Two methods of RNAseq analysis were used. For neuronal cultures, filtering of low-express genes and trimmed mean of M-values (TMM) normalization were performed before differential gene expression analysis with the limma-voom pipeline ^84^. For microglial sequencing data, manual filtering was performed to remove low-expressed genes and retain only protein-coding genes. We identified large differences in library sizes between samples in our microglial RNA-seq. To account for this variation, we followed the DESeq2 analysis pipeline, which normalizes data by calculating sample-specific size factors to adjust for sequencing depth and library composition ^85^. Biological replicates were plated and grown in individual wells, and separate libraries were prepared and sequenced per replicate. Thus, these were considered biological replicates and therefore not combined using Deseq2 tools, such as collapseReplicates, as this is designed for technical replicates only. DESeq2 normalizations and differential gene expression analysis were performed, and finally, gene set enrichment analysis (GSEA) was performed using clusterProfiler ^86^ using MSigDB gene set database with annotations ^87–89^. Statistical significance was determined as a p-value of 0.05 or below, adjusted with the Benjamini-Hochberg multiple corrections.

The spatial transcriptomic data were initially handled using the Nanostring/Bruker GeoMx NGS pipeline, converting FASTQ files to digital count conversion files. Segments were excluded based on low raw read threshold, percentage aligned reads, and sequence saturation in R. Filtering was performed to separate distinct cell types and regions. Using the StandR package ^90^, gene-level quality control and data cleaning were performed through low-expression gene filtering with minimum counts calculated according to a 2X multiplication of mean negative probe counts, and stringent sample fraction thresholds as parameters for removal. TMM normalizations were performed using the StandR geomxNorm function, and thereafter, slide-specific variation was investigated. PCA was used to confirm the source of further unexplained sample variation, and to determine the top removed genes in the data (Supplementary Figure 7). Consequently, remove unwanted variation 4-step (RUV4) was selected for batch corrections. For further data quality assessment, AOI nuclei counts were inspected, indicating that the negative probe signal does not increase as area increases as expected. Thus, to ensure sufficient quality of RNA, negative probe counts were assessed alongside manual inspection of the data. Thus, AOI nuclei counts were not used in the data analysis process to reduce the risk of potential bias (Supplementary Figure 8). Final filtering was performed before differential gene expression analysis with limma-voom ^84^.

Thereafter, GSEA was performed using Gene Ontology Biological Process (GOBP) terms. In addition, a targeted GSEA analysis was performed to identify pathways exclusively associated with the relevant tissue and cell type. This selection was performed by filtering gene-sets for titles containing common terms associated with the brain and microglia. Gene-sets included were KEGG, GOBP, Wikipathways, and REACTOME, and these were filtered for the following terms: ‘microglia’, ‘neuro’, ‘inflam’, ‘immune response’, ‘cytokine’, ‘glia’, ‘brain’, ‘synap’, ‘myelin’, and ‘nervous’.

### Rank based cell type enrichment analysis

Cell-type (in this case, microglia sub-type / state) enrichment analysis was performed using two curated cell-type marker gene sets. Rank based cell-type enrichment analysis was performed using fast gene set enrichment analysis (FGSEA) ^91^. Genes were ranked according to differential expression statistics (log fold-change), and duplicate gene symbols were collapsed by retaining the entry with the largest absolute fold-change. Cell-type marker sets were tested for enrichment across the full ranked transcriptome using FGSEA, allowing detection of coordinated transcriptional shifts without applying hard DEG thresholds. For each cell type, normalized enrichment scores (NES), adjusted p-values, enrichment scores (ES), and leading-edge genes driving the enrichment signal were obtained. Multiple testing correction was performed using the Benjamini–Hochberg method.

Plots were generated including a vertical dashed line indicating the significance threshold corresponding to an adjusted p-value (FDR) of 0.05. Points to the right of this threshold represent statistically significant enrichments. Enrichments observed across multiple related MG states may reflect shared transcriptional programs between closely related microglial subpopulations.

Reference datasets used in FGSE analysis include a human iPSC-derived microglia dataset in which cells were exposed to brain substrates ^15^ and was accessed at https://doi.org/10.1038/s41590-023-01558-2 in supplementary table 2. The reference dataset used in FGSE analysis of our *anx/anx* murine data was a transcriptomic taxonomy atlas of microglia from murine models of several neurodegenerative and inflammatory disorders and was accessed at https://doi.org/10.1038/s41590-026-02472-z from a processed Seurat object available at https://doi.org/10.5281/zenodo.16938034

## Code availability

All analyses were done using the R programming language ^92^. More information on R environment, packages, and scripts, on https://github.com/The-Neurobiology-of-Anorexia-Nervosa/Aberrant-microglial-responses-shape-hypothalamic-circuits-in-anorexia-nervosa.

## Data availability

Transcriptomic data will be made available for download from the NCBI Gene Expression Omnibus.

## Author contributions

The hypothesis was generated by JX, CS and IAKN. JX, EE, KZ, FO, SS, MS, CS, BE, CC, EW, LS, TH, and IAKN designed the study. JX, EE, TF, CC, BE, KZ, MC, LS, TH, and IAKN performed experiments and/or conducted analyses. All authors interpreted data. The manuscript was drafted by JX, EE, and IAKN, reviewed for intellectual content and approved by all authors.

## Acknowledgements

We are grateful for the help from Marie Carp with recruiting the study participants, Studiebehandlingsenheten at the Karolinska Hospital with sampling of study participants, as well as the discussions and advices regarding spatial transcriptomic data analysis with Dr. Roman Romanov (Medical University of Vienna), and regarding bioinformatics with Dr. Aditya Sing and the Swedish Bioinformatics Advisory Program (National Bioinformatics Infrastructure [NBIS], SciLife Labs). KIGene core facility at Karolinska Institutet/Karolinska Hospital provided the platform enabling spatial transcriptomics. The RNA sequencing data analysis was enabled by resources provided by the National Academic Infrastructure for Supercomputing in Sweden (NAISS), partially funded by the Swedish Research Council through grant agreement no. 2022-06725. Lastly, we wish to thank the study participants for their invaluable contribution to this work.

