## Supplementary Figures and Tables for "Aberrant microglial responses shape hypothalamic circuits in anorexia nervosa"

### Supplementary Information

Jingjing Xu[1,2*], Emmy Erskine[1,2#*], Barbara Eramo[1,2], Karin Zimmer[1,2], Chiara Camoglio[1,2], Mridul Chaudhary[3], Tianxu Feng[1,2], Funda Orhan[3], Elisabeth Welch[4,5], Lars Selander[1,2], Catharina Lavebratt[1,2],Tomas Hökfelt[6], Martin Schalling[1,2], Samudyata Samudyata[3], Carl M. Sellgren[3,6,7§], Ida AK Nilsson[1,2,8#§]

*[1]Department of Molecular Medicine and Surgery, Karolinska Institutet, Stockholm, Sweden*

*[2]Center for Molecular Medicine, Karolinska University Hospital, Stockholm, Sweden*

*[3]Department of Physiology and Pharmacology, Karolinska Institutet, Stockholm, Sweden*

*[4]Department of Clinical Neuroscience, Karolinska Institutet, Stockholm, Sweden.*

*[5]Department of Women's and Children's Health, Uppsala University, Akademiska sjukhuset, Uppsala, Sweden.*

*[6]Department of Neuroscience, Karolinska Institutet, Stockholm, Sweden*

*[7]Center for Psychiatry Research, Department of Clinical Neuroscience, Karolinska Institutet*

*[8]Stockholm Health Care Services, Stockholm County Council, Stockholm, Sweden.*

*[9]Center for Eating Disorders Innovation, Karolinska Institutet, Stockholm, Sweden*

** shared first*

*§ shared last*

**Table of contents: Supplementary materials**


Supplementary Table 1 (attached)
CNVs

Supplementary Table 2 (below)
Antisera

Supplementary Table 3 (below)
Primer Sequences

Supplementary Table 4 (attached)
Phagocytosis

Supplementary Table 5 (attached)
DEGs

Supplementary Table 6 (attached)
GSEA

Supplementary Table 7 (attached)
FGSEA

Supplementary Table 8
IMARIS quantification

Supplementary Figure 1 (below)
iPSC stainings

Supplementary Figure 2 (below)
Cell counts, synaptosome yield

Supplementary Figure 3 (below)

Overview of *in vitro* differentiations and transcriptomic analysis

Supplementary Figure 4( below)

qPCR iMGs GLP1R

Supplementary Figure 5 (below)

Transcriptomic analysis of AN microglia upon GLP1 stimulation and after phagocytosis of hypothalamic and cortical synaptosomes

Supplementary Figure 6 (below)

DEGs of hypothalamic neurons vs cortical neurons

Supplementary Figure 7 (below)

Spatial transcriptomics method

Supplementary Figure 8 (below)

GeoMX AOI nuclei count investigation

Supplementary figure 9 (below)

Spatial transcriptomics of *anx/anx* mouse

**Supplementary Table 2**. List of antibodies used in the study

| Antibody | Company | Cat. Number | Species | Concentration |
|---|---|---|---|---|
| IBA1 Polyclonal | Wako | 019-19741 | Rabbit | 1:200 |
| SYN½ Polyclonal | Synaptic Systems | 106 002 | Rabbit | 1:3000 |
| IBA1 Polyclonal | Novus Biolabs | NB100-1028 | Goat | 1:200 |
| α-MSH | Millipore | ab5087 | Sheep | 1:200 |
| MAP2 | Abcam | ab5392 | Chicken | 1:1000 |
| TBR1 | Abcam | ab31940 | Mouse | 1:100 |
| VGAT | Synaptic Systems | 131,011 | Mouse | 1:500 |
| P2Y12 | Invitrogen | 702516 | Rabbit | 1:200 |

**Supplementary Table 3**. Primer sequences

| Primer sequences | Forward | Reverse | Final concentration |
|---|---|---|---|
| GLP1R | CCTCCTGCCACAGACTTGTT | GCTGACATTCACGAACGAGC | 500nM |
| ACTB | GATGCAGAAGGAGATCACTGC | ATACTCCTGCTTGCTGATCCA | 500nM |

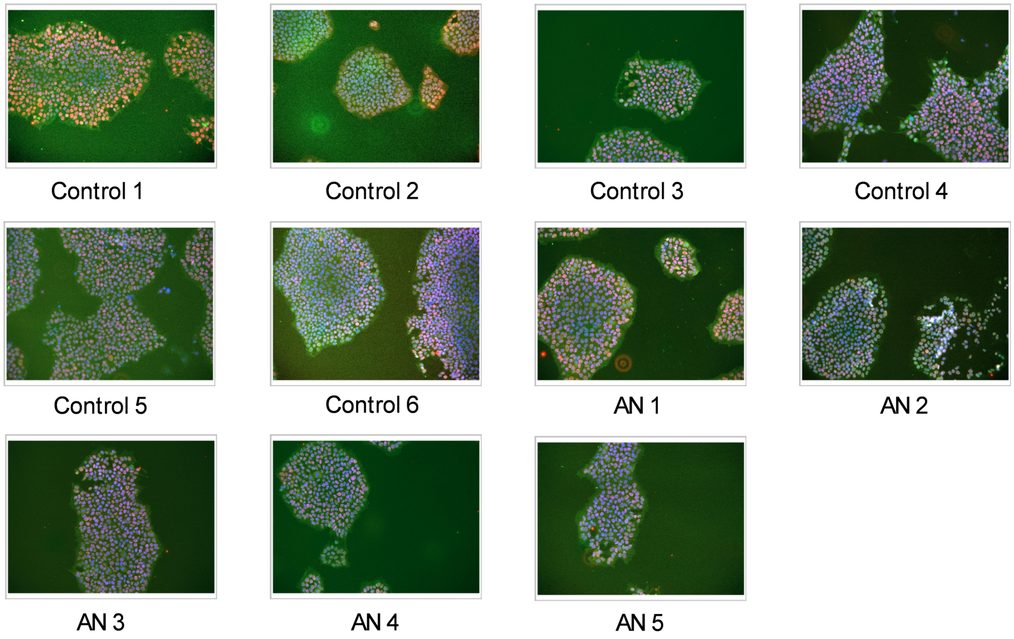


**Supplementary Figure 1: Quality control of iPSC lines**

Representative ICC images of all iPSC lines stained for pluripotency markers NANOG (green) and OCT4 (red), in combination with DAPI (blue). AN, anorexia nervosa; iPSC, inducible pluripotent stem cells; ICC, immunocytochemistry; NANOG, homeobox protein NANOG; OCT4, Octamer binding transcription factor 4.

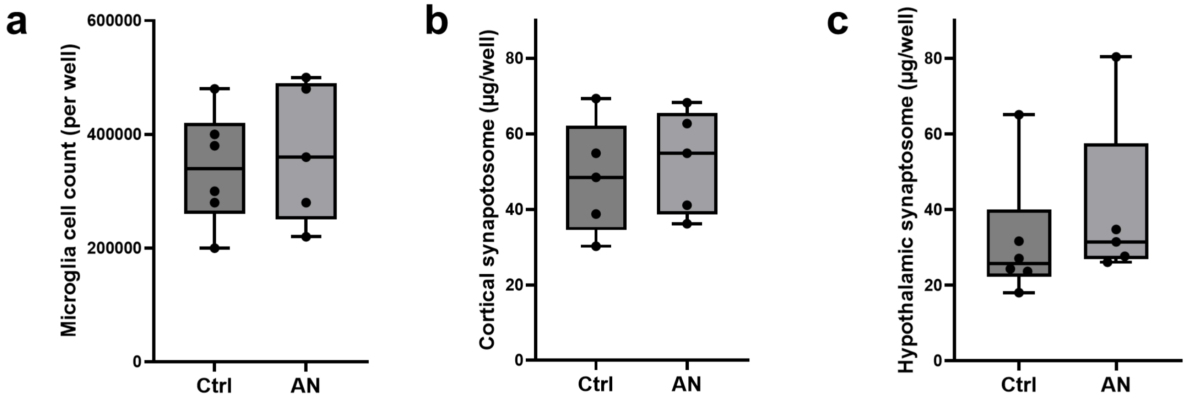


**Supplementary Figure 2: quantification of microglia and synaptosomes**

**(a)** Microglial cell yield per well of a 6-well plate. Each data point represents the mean cell count for each cell line from three independent experiments. **(b, c)** Yields of cortical and hypothalamic synaptosomes isolated per well of a 6-well plate. Each data point represents the average yield of each line from two independent experiments. Ctrl, control; AN, anorexia nervosa.

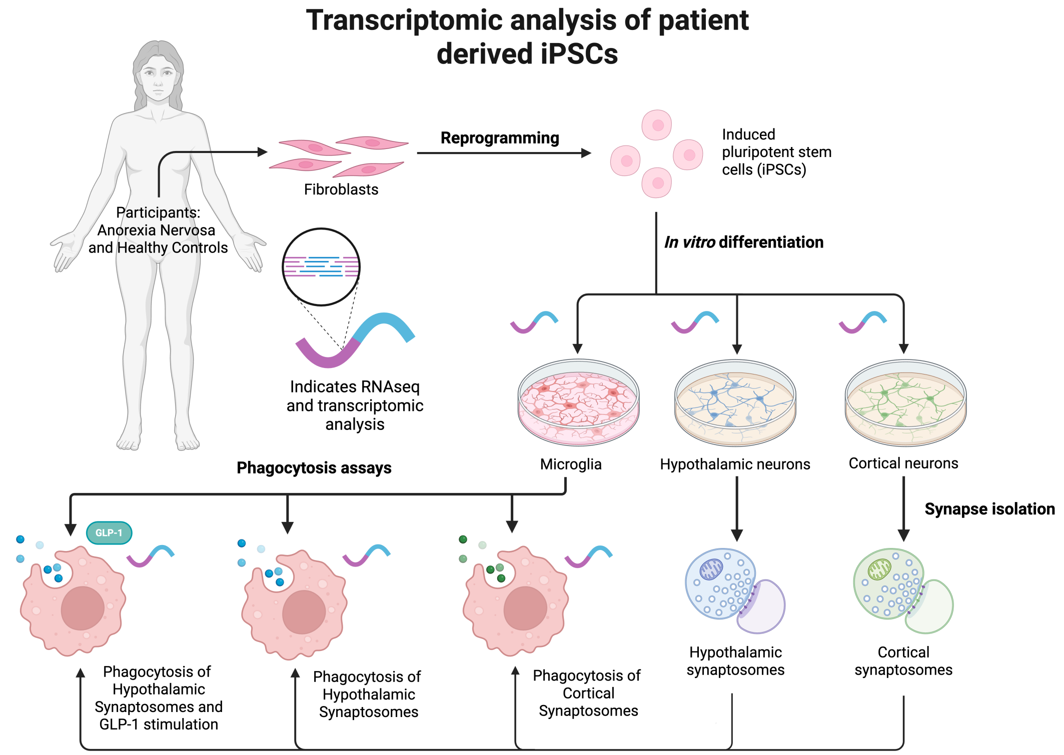


**Supplementary Figure 3: Overview of transcriptomic analysis of patient derived iPSCs**

Schematic overview of patient samples, differentiated microglia, hypothalamic and cortical neurons, stimulations and phagocytosis assays. AN, Anorexia nervosa; HC, Healthy Control; GLP1, glucagon-like peptide 1. Created in BioRender. Erskine, E. (2026) https://BioRender.com/13cp0lq.

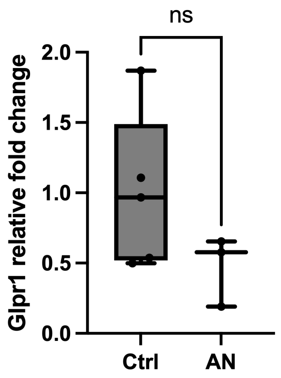


**Supplementary Figure 4: GLP1R qPCR**

Relative expression of GLP1R (p = 0.1834) in microglia determined by qPCR. Ctrl, control; AN, anorexia nervosa; GLPR1, glucagon-like peptide receptor 1; qPCR, quantitative polymerase chain reaction; ns, non significant.

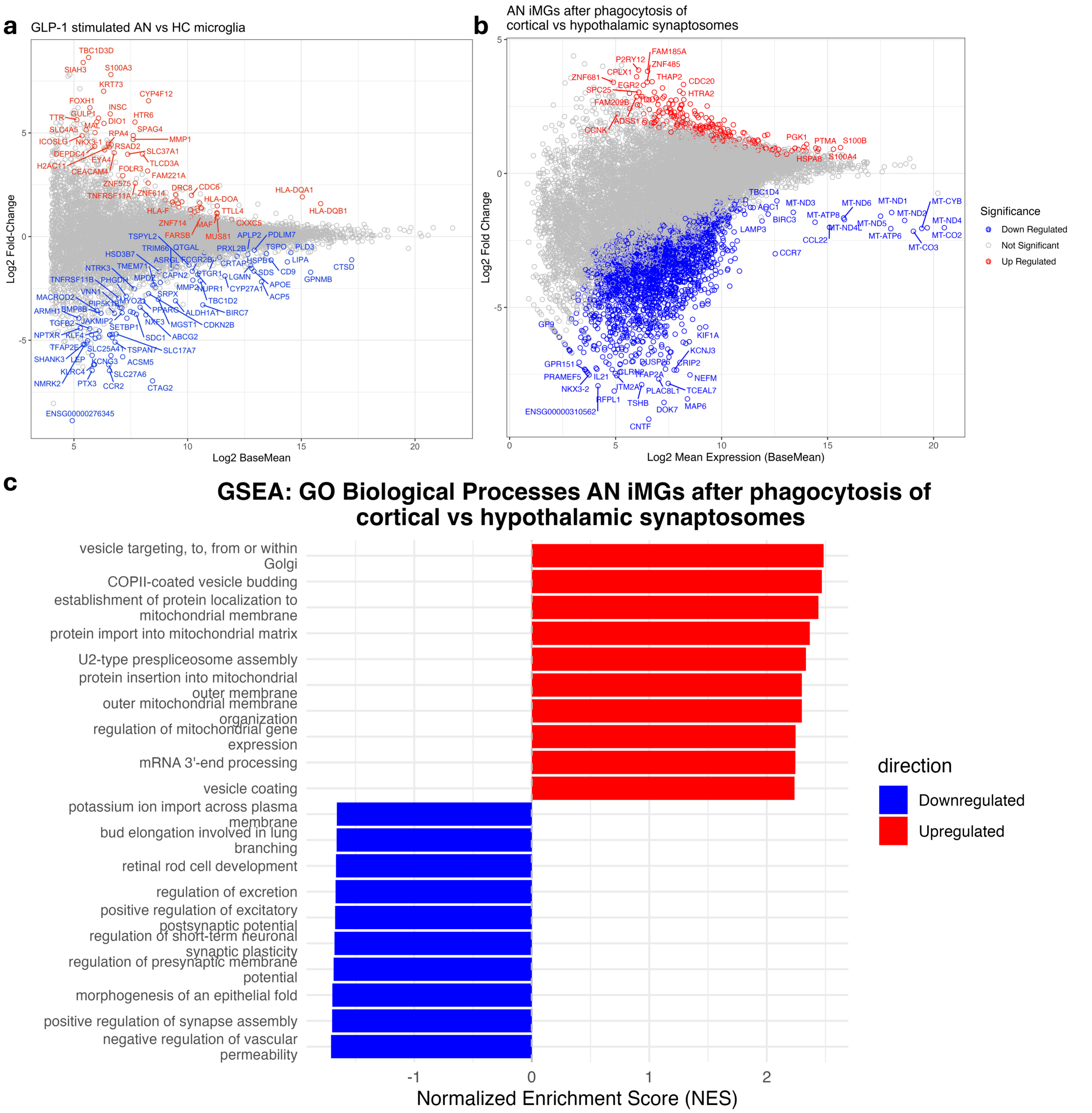


**Supplementary figure 5: Transcriptomic data analysis of AN microglia after stimulation with GLP-1 and after phagocytosis of cortical vs hypothalamic synaptosomes**

(a) DEG analysis of AN microglia upon GLP-1 treatment. (b) DEGs of AN microglia after phagocytosis of cortical vs hypothalamic synaptosomes (b) GSEA GOBP terms in AN microglia after phagocytosis of cortical vs hypothalamic synaptosomes. Red/blue corresponds to upregulated/ downregulated genes ($p < 0.05$). DEG, differentially expressed gene; AN, Anorexia nervosa; iMG, induced microglia; GSEA gene set enrichment analysis; GOBP, gene ontology biological processes; HC, Healthy Control; GLP1, glucagon-like peptide 1.

a

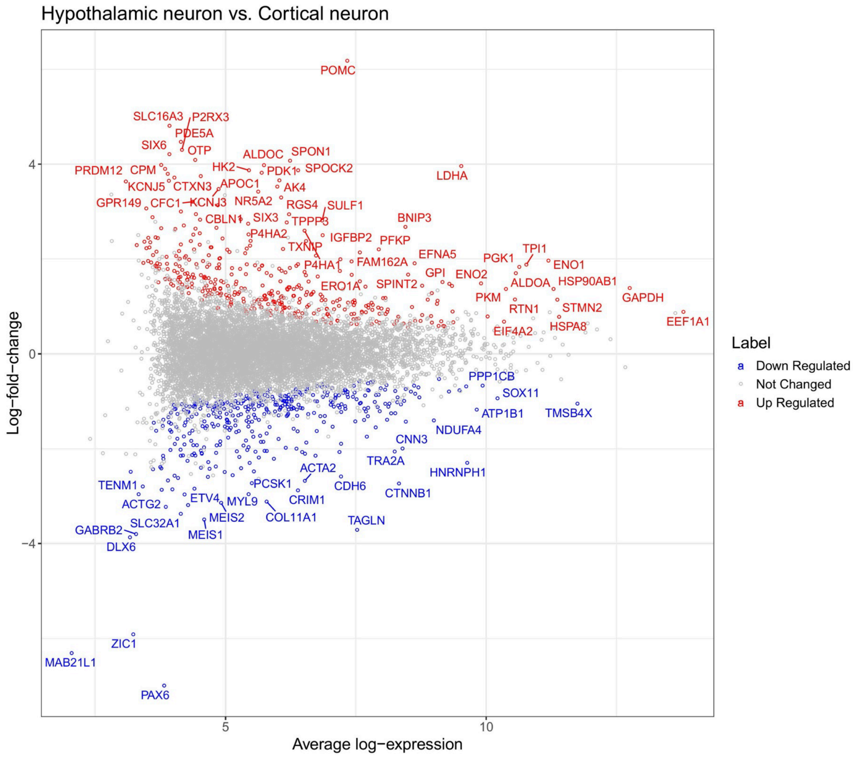


b

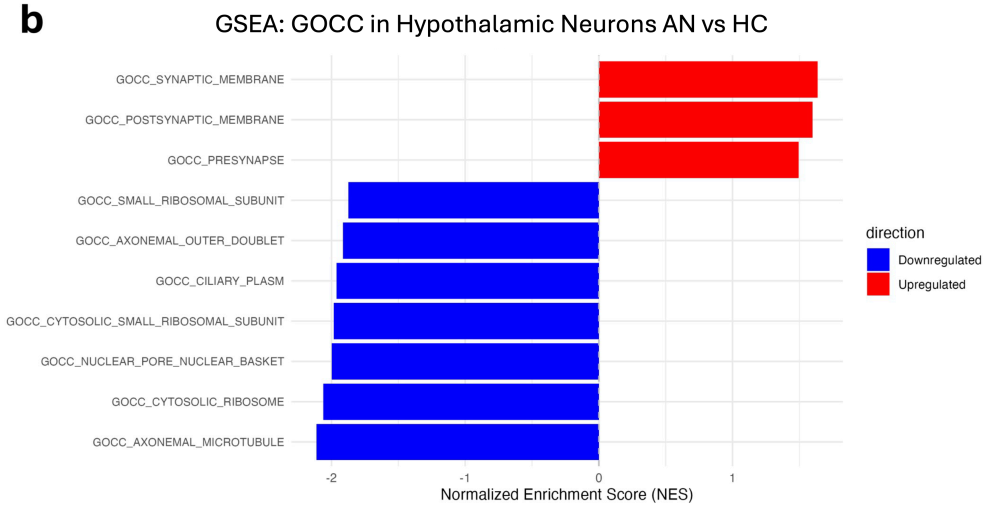


**Supplementary Figure 6: DEGs of hypothalamic neurons vs cortical neurons**

DEG analysis of hypothalamic neurons compared to cortical neurons. Red/blue corresponds to upregulated/ downregulated genes ($p < 0.05$). DEG, differentially expressed gene; GSEA, gene set enrichment analysis; GOBP, gene ontology biological processes

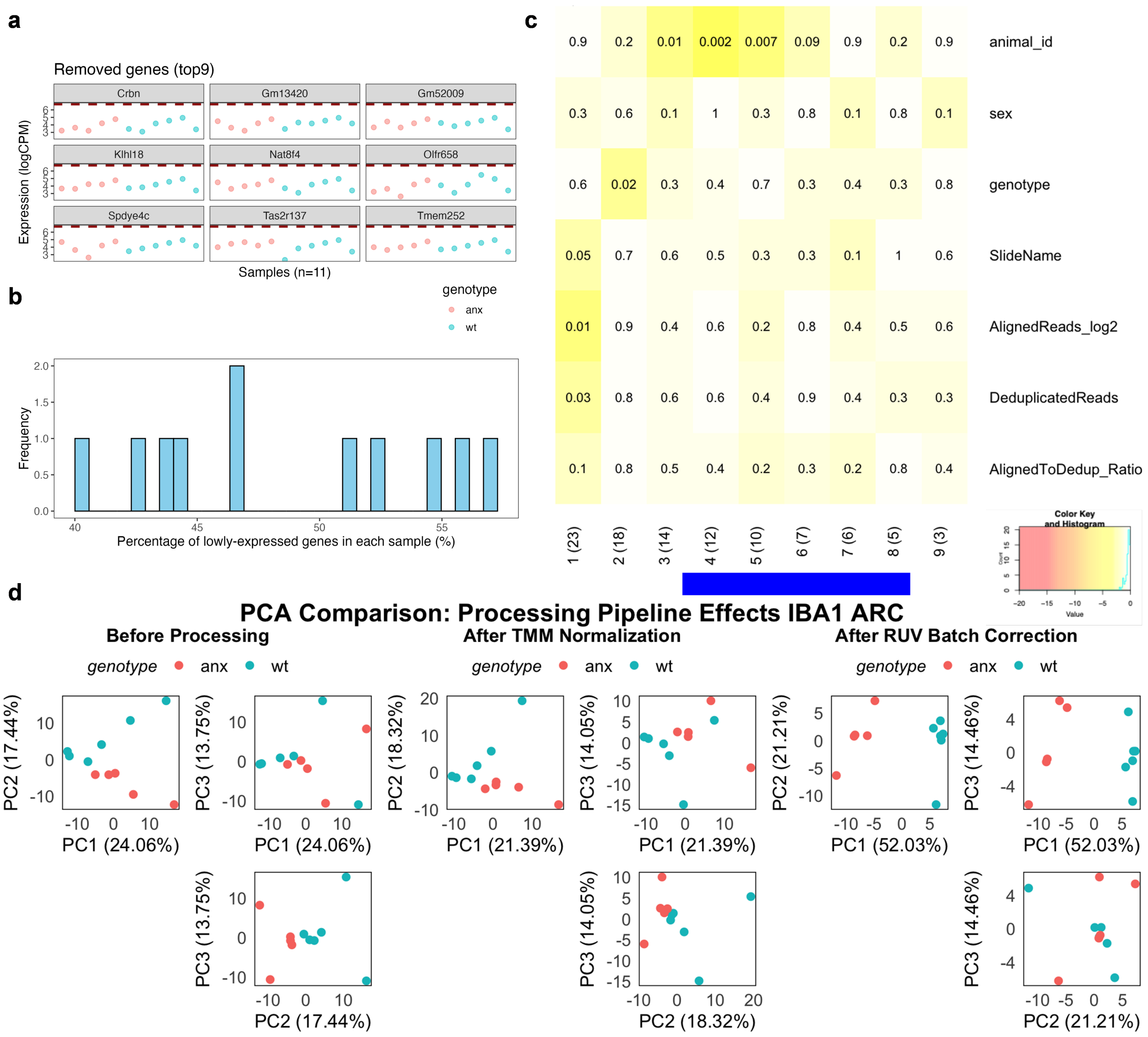


**Supplementary figure 7: Spatial transcriptomic data analysis quality control with variance testing and RUV4 batch corrections illustrated with the IBA1 ARC dataset**

a) top genes removed during low expression gene filtering b) percentage of lowly expressed genes in samples c) Principal component heatmap assessing explained and unexplained variance d) PCA plots assessing explained variance before and after normalizations and batch corrections with RUV4. WT, wild-type; PCA, principle component analysis; IBA1, ionized calcium binding adaptor molecule 1; ARC, arcuate hypothalamus.

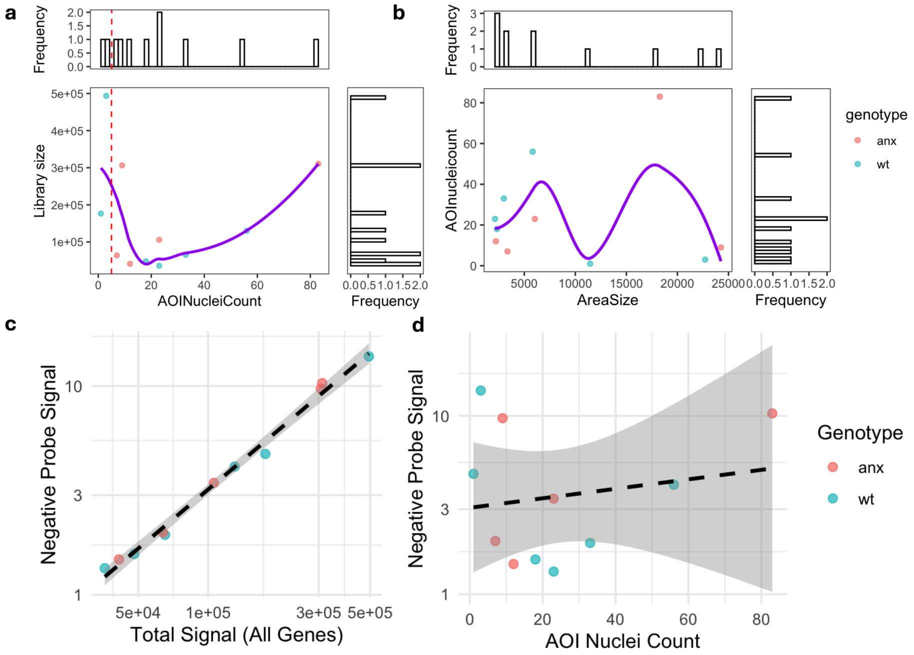


**Supplementary Figure 8: Quality control of AOI nuclei counts in GeoMX spatial transcriptomic data**.

a) library size against AOINucleiCount plot b) AOINucleiCount against area size c) negative probe signal against total signal d) negative probe signal against AOI nuclei count. In all plots, anx is pink and WT is blue. AOI; area of interest; WT, wild-type.

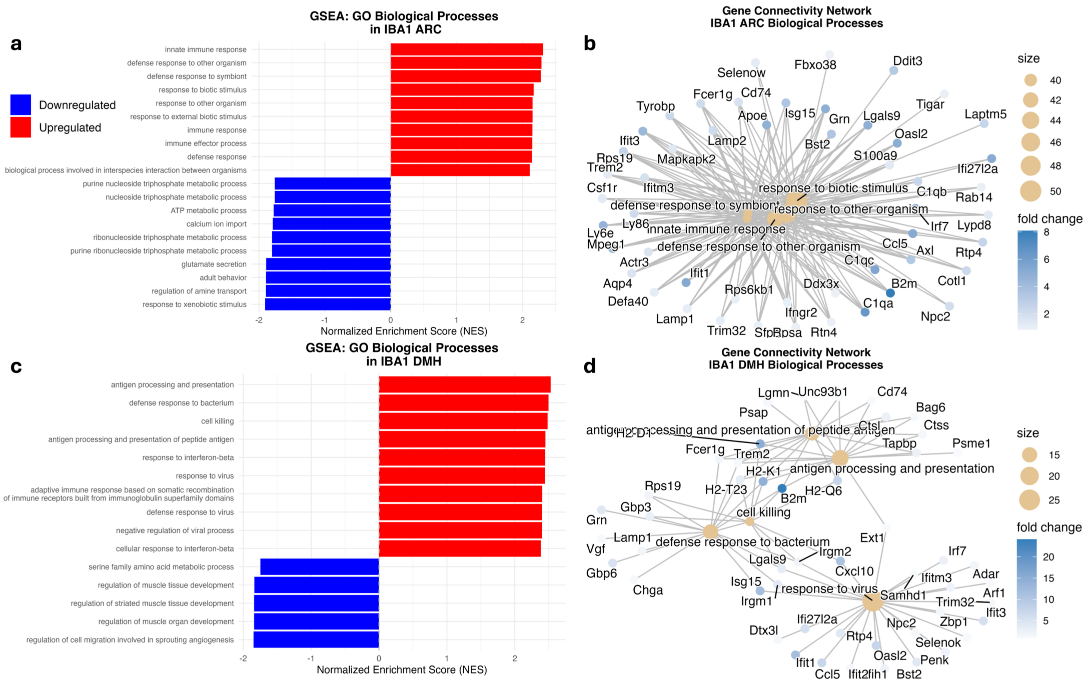


**Supplementary figure 9: Spatial transcriptomic data analysis *anx/anx* mouse, GSEA, and Network analysis**

a) GSEA GOBP terms in IBA1 ARC (p values) b) Cnet network analysis plots of IBA1 ARC GOBP positive terms c) GSEA GOBP terms in IBA1 DMH (p values) d) Cnet network analysis plots of IBA1 DMH GOBP positive terms. Colour of GSEA analysis represents the up or downregulation (upregulated in red and downregulated in blue). In Cnet plots blue colour gradient indicates fold change and the size of the dots represents the size of the pathway. IBA1, Ionized calcium binding adaptor molecule 1; ARC, arcuate hypothalamus; DMH; dorsomedial hypothalamus GSEA gene set enrichment analysis; GOBP, gene ontology biological processes; Cnet category gene network.
